# SOCS3-mediated activation of p53-p21-NRF2 axis and cellular adaptation to oxidative stress in SOCS1-deficient hepatocellular carcinoma

**DOI:** 10.1101/2021.10.21.465149

**Authors:** Md Gulam Musawwir Khan, Nadia Boufaied, Mehdi Yeganeh, Amit Ghosh, Rajani Kandhi, Rezvan Bagheri, Stephanie Petkiewicz, Ankur Sharma, Akihiko Yoshimura, Gerardo Ferbeyre, David P. Labbé, Sheela Ramanathan, Subburaj Ilangumaran

## Abstract

*SOCS1* and *SOCS3* genes, frequently repressed in hepatocellular carcinoma (HCC), function as tumor suppressors in hepatocytes. However, TCGA transcriptomic data revealed that *SOCS1-low/SOCS3-high* specimens displayed more aggressive HCC than *SOCS1-low/SOCS3-low* cases. We show that hepatocyte-specific *Socs1-*deficient livers upregulate *Socs3* expression following genotoxic stress. Whereas deletion of *Socs1* or *Socs3* increased HCC susceptibility, ablation of both genes attenuated HCC growth. SOCS3 promotes p53 activation in SOCS1-deficient livers, leading to increased expression of CDKN1A (p21^WAF1/CIP1^), which coincides with elevated expression and transcriptional activity of NRF2. Deleting *Cdkn1a* in SOCS1-deficient livers diminished NRF2 activation, oxidative stress and HCC progression. Elevated *CDKN1A* expression and enrichment of antioxidant response genes also characterized *SOCS1-low/SOCS3-high* HCC. SOCS1 expression in HCC cell lines reduced oxidative stress, p21 expression and NRF2 activation. Our findings demonstrate that SOCS1 controls the oncogenic potential of SOCS3-driven p53-p21-NRF2 axis and suggest that NRF2-mediated antioxidant response represents a drug target in SOCS1-deficient HCC.

## Introduction

SOCS1 and SOCS3 generally function as negative feedback regulators of cytokine, growth factor and Toll-like receptor signaling and can also activate the tumor suppressor p53 ^1–6^. A tumor suppressor function for SOCS1 in the liver was proposed as the *SOCS1* gene is frequently repressed by epigenetic mechanisms in hepatocellular carcinoma (HCC) ^7, 8^. Supporting this notion, mice lacking SOCS1 in hepatocytes show increased susceptibility to diethylnitrosamine (DEN)-induced HCC ^9, 10^. *SOCS3* is also repressed in HCC albeit to a lower extent than *SOCS1*, and hepatocyte-specific SOCS3-deficient mice show increased DEN-induced HCC ^11, 12^. Even though SOCS1 and SOCS3 share close structural similarity ^13^, increased HCC development in mice lacking either SOCS1 or SOCS3 indicate their non-overlapping tumor suppressor functions.

Mice lacking either *Socs1* or *Socs3* show accelerated liver regeneration, indicating that SOCS1 and SOCS3 play key roles in regulating hepatocyte proliferation ^12–14^. During liver regeneration, SOCS3 deficiency in hepatocytes deregulates IL-6 signaling, while SOCS1 deficiency potentiates HGF signaling ^12, 14, 15^. As cytokines and growth factors that facilitate physiological hepatocyte proliferation also drive hepatocarcinogenesis ^16^, SOCS1 and SOCS3 likely mediate their tumor suppressor functions, at least partly, via attenuating HGF and IL-6 signaling, respectively. Studies on oncogene-induced senescence implicated SOCS1 in activating p53 ^4, 6^, suggesting that SOCS1 deficiency may compromise p53-mediated tumor suppression. However, SOCS1 deficiency did not diminish the induction of p53 target genes in the liver following genotoxic stress ^10^. In contrast, SOCS1 deficiency increased the expression of the p53 target gene *Cdkn1a* and SOCS1 expression in hepatocytes reduced the stability of CDKN1A (p21) protein ^10^. As a cyclin-dependent kinase (CDK) inhibitor, CDKN1A generally functions as a tumor suppressor ^17^. Paradoxically, many tumors including HCC show elevated CDKN1A expression that correlates with high malignancy, poor prognosis and drug resistance ^17, 18^. Deregulated growth factor signaling in SOCS1-deficient hepatocytes promotes AKT-mediated phosphorylation of CDKN1A, retaining it in the cytosol, where it can exert pro-tumorigenic effects ^10, 17^. CDKN1A also interacts with nuclear factor erythroid 2-related factor 2 (NFE2L2 or NRF2), leading to NRF2-mediated antioxidant response that is exploited by cancers to tolerate the increased oxidative stress associated with neoplastic growth ^19–21^. These reports suggest that the loss of SOCS1-dependent regulation could render CDKN1A oncogenic, however genetic evidence for this hypothesis is lacking. Moreover, in contrast to SOCS1, SOCS3 was shown to promote the expression of CDKN1A ^5, 22^, and the impact of this pathway on HCC is not known.

Despite tremendous advances in understanding the molecular pathways, oncogenes and tumor suppressors, and extensive genetic profiling of human cancers, targeted molecular therapeutics has made only a moderate impact on disease outcome and the survival rate in HCC ^23, 24^. Combining molecular therapies with immunotherapy or targeting cancer metabolites are proposed to improve this scenario ^23, 25^. However, it is becoming increasingly clear that cellular adaptation during oncogenesis can alter normal functions of many proteins such as CDKN1A and NRF2 tumor suppressors ^17, 21^, which may exert completely opposite effects depending on the molecular context of cancers. Understanding the molecular basis of the context-dependent alterations in protein functions, aided by the genetic signature of individual cancers, though daunting, will be crucial to realize the full potential of targeted molecular therapies. Towards this direction, we sought to dissect the impact of SOCS1 and SOCS3, alone and together, on HCC and their role in modulating CDKN1A using data from human patients and mouse genetic models.

## Results

### *SOCS1-low/SOCS3-high* HCC show poor survival

Even though mouse genetic studies have shown that loss of either *Socs1* or *Socs3* increases susceptibility to HCC ^1, 9, 12^, the Cancer Genome Atlas on liver HCC (TCGA-LIHC) transcriptomic data ^24^ predicted poor progression-free survival only for low *SOCS1* but not for low *SOCS3* expression (**Figure 1A**). To determine whether a diminished expression of both *SOCS1* and *SOCS3* would increase disease severity, we stratified the HCC cases based on *SOCS1* (low, high) as well as *SOCS3* (low, high) expression (**Figure 1B**). Strikingly, Kaplan-Meir curves revealed that *SOCS1-low/SOCS3- high* tumors displayed the faster disease progression than *SOCS1-low/SOCS3-low* cases (**Figure 1C**). This feature was also strengthened by a trend towards a higher hazard ratio (HR) for the *SOCS1- low/SOCS3-high* group in univariate analysis (HR = 1.80, 95% Confidence Interval (CI) 0.99-3.3; *p*=0.053; **Figure 1D**). These findings indicated an adverse impact of high *SOCS3* expression on tumor progression in *SOCS1-low* HCC patients.

**Figure 1.**
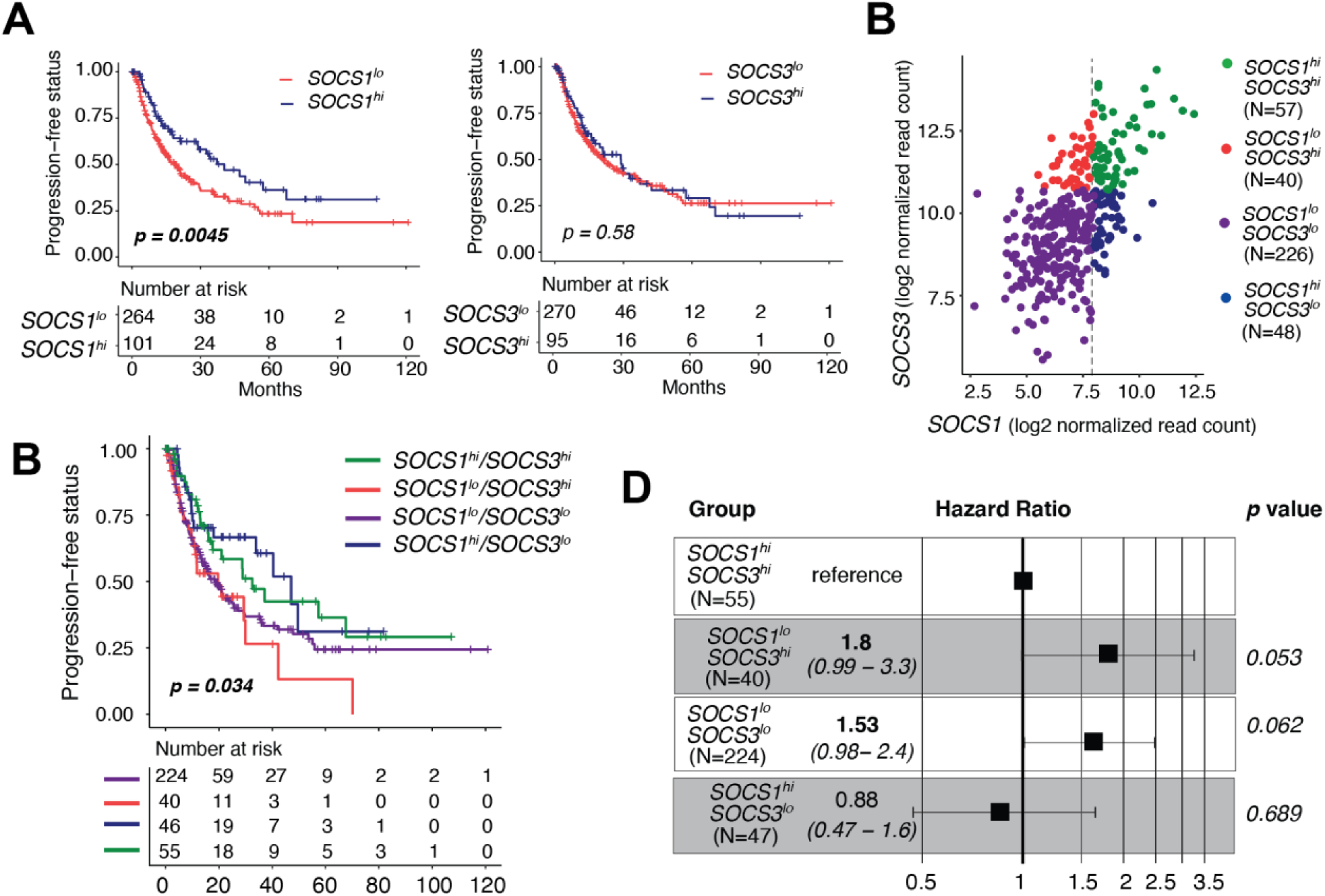
High *SOCS3* expression in *SOCS1-low* HCC cases in the TCGA-LIHC cohort predicts faster disease progression. (A) Low SOCS1 but not SOCS3 predicts poor progression-free survival in the TCGA-LIHC cohort. (B) Segregation of the TCGA-LIHC dataset into *SOCS1-high/SOCS3-high*, *SOCS1-low/SOCS3-high*, *SOCS1-low/SOCS3-low* and *SOCS1-high/SOCS3-low* groups based on z-score. (C) Poor progression-free survival for the *SOCS1-low/SOCS3-high* group. (D) increased hazard ratio for the *SOCS1-low/SOCS3-high* group.

### SOCS3 promotes HCC progression in SOCS1-deficient livers

To determine how SOCS3 impacted HCC development in the absence of SOCS1, we first evaluated hepatic *Socs1* and *Socs3* gene expression in mice lacking either SOCS1 or SOCS3 in hepatocytes following administration of diethylnitrosamine (DEN). *Socs3* was upregulated nearly 16-fold in *Socs1^fl/fl^Alb-Cre* mice, whereas *Socs1* induction in *Socs3^fl/fl^Alb-Cre* mice was comparable to that of control mice (**Figure 2A**), suggesting a compensatory induction of *Socs3* in SOCS1-deficient hepatocytes but not vice versa. We generated mice lacking both SOCS1 and SOCS3 in hepatocytes and examined their susceptibility to developing HCC ten months after the administration of DEN at two weeks of age. Whereas *Socs1^fl/fl^Alb-Cre* and *Socs3^fl/fl^Alb-Cre* mice developed HCC with 100% penetrance and showed significantly greater number and size of tumor nodules than *Socs1^fl/fl^* and *Socs3^fl/fl^* controls (**Figure 2B,C**, **Figure S1**), simultaneous loss of SOCS1 and SOCS3 did not result in aggressive HCC. In fact, *Socs1^fl/fl^Socs3^fl/fl^Alb-Cre* mice showed significantly reduced tumor volume compared to *Socs1^fl/fl^Alb-Cre* mice, even though the incidence and the number of tumor nodules remained comparable (**Figure 2B, C**). The tumor nodules from mice lacking either SOCS1 or SOCS3 harbored significantly more Ki67-positive cells than those from control mice, whereas *Socs1^fl/fl^Socs3^fl/fl^Alb-Cre* mice showed fewer proliferating cells (**Figure 2D,E**, **Figure S2**). These data indicated that genotoxic stress in SOCS1-deficient hepatocytes causes compensatory SOCS3 induction, and that SOCS3 expression is necessary to sustain increased tumor growth in SOCS1- deficient livers.

**Figure 2.**
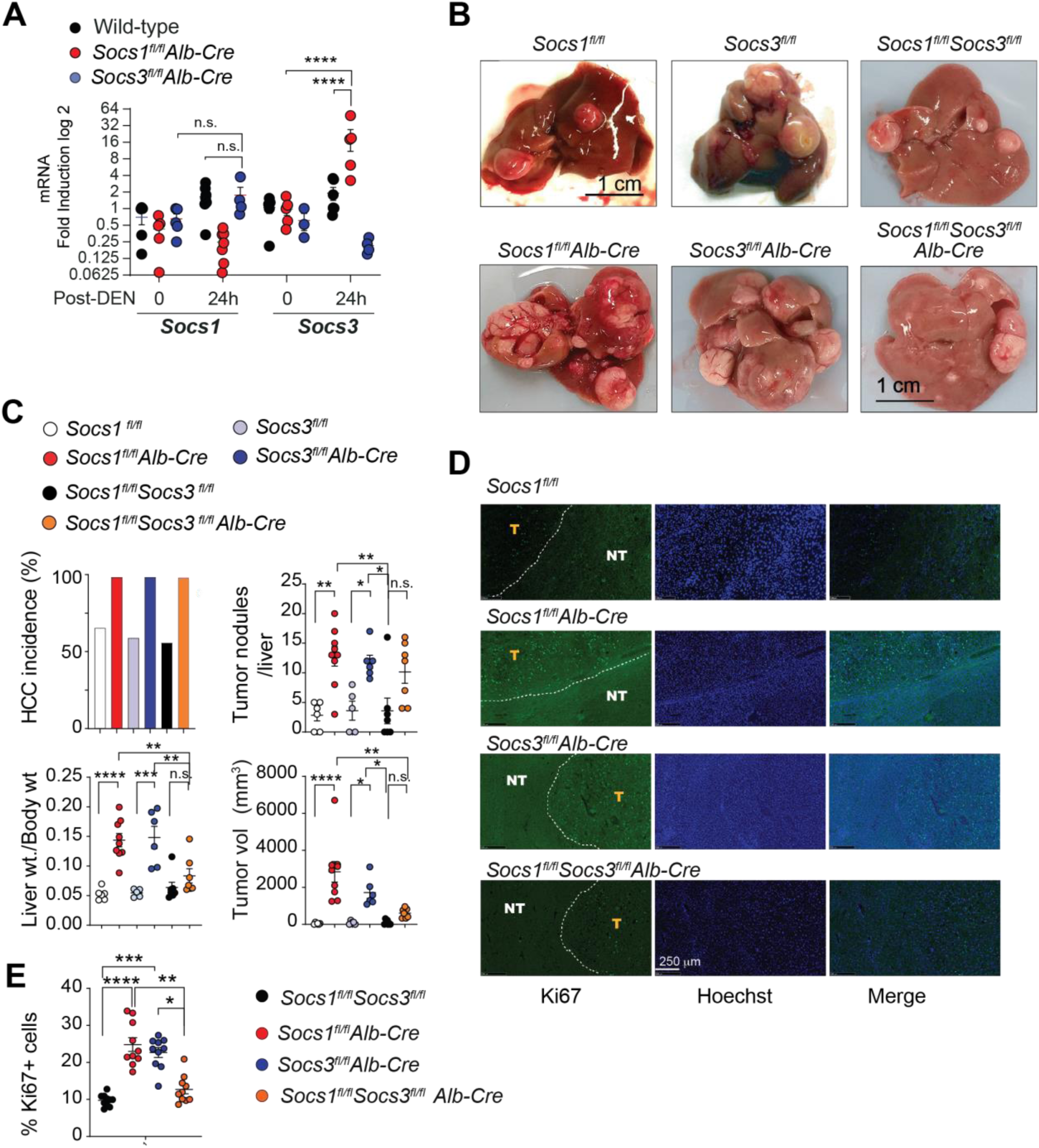
Compensatory SOCS3 induction in SOCS1-deficient hepatocytes promotes HCC growth. (A) *Socs1* and *Socs3* gene expression in the livers of hepatocyte-specific SOCS1- or SOCS3- deficient mice following DEN treatment (100 mg/Kg bodyweight; n=5 mice/group). Mean ± SEM. (B) HCC development in mice lacking SOCS1, SOCS3 or both in hepatocytes 10 months after injection of DEN (25 mg/Kg bodyweight) at two weeks of age. Representative macroscopic liver images of n=5-9 mice per group are shown. (C) Cumulative data on incidence rate, number of visible tumors and tumor volume in mice groups shon in (B). (D) Ki67 immunostaining in DEN-induced HCC in the mice groups shown in (B). Representative sections from ≥3 mice/group shown. Tumor nodules (T) are demarcated from surrounding normal tissue (NT). (E) Quantification of Ki67+ nuclei over Hoechst-stained nuclei from 10 randomly selected areas from 3 mice per group. Statistics: One-way (C, E) or two-way (A) ANOVA with Tukey’s multiple comparison test; \**p*<0.05, \*\**p*<0.01, \*\*\**p*<0.001, \*\*\*\**p*<0.0001.

### SOCS3 promotes oncogenesis via p53-dependent CDKN1A induction

SOCS1 deficiency increases the expression of CDKN1A and promotes its oncogenic potential in a murine HCC model ^10^. CDKN1A is a transcriptional target of p53, which can be activated by both SOCS1 and SOCS3 ^4, 5, 17^. Therefore, we examined whether SOCS3 modulated DEN-induced *Cdkn1a* expression in *Socs1^fl/fl^Alb-Cre* mice. *Cdkn1a* was induced in *Socs1^fl/fl^Alb-Cre* mice several hundred-fold more strongly than in control mice and to a lesser extent in *Socs3^fl/fl^Alb-Cre* mice (**Figure 3A**). However, the combined loss of SOCS1 and SOCS3 completely abrogated *Cdkn1a* induction, indicating that SOCS3 is crucial for *Cdkn1a* induction in SOCS1-deficient hepatocytes (**Figure 3A**).

**Figure 3.**
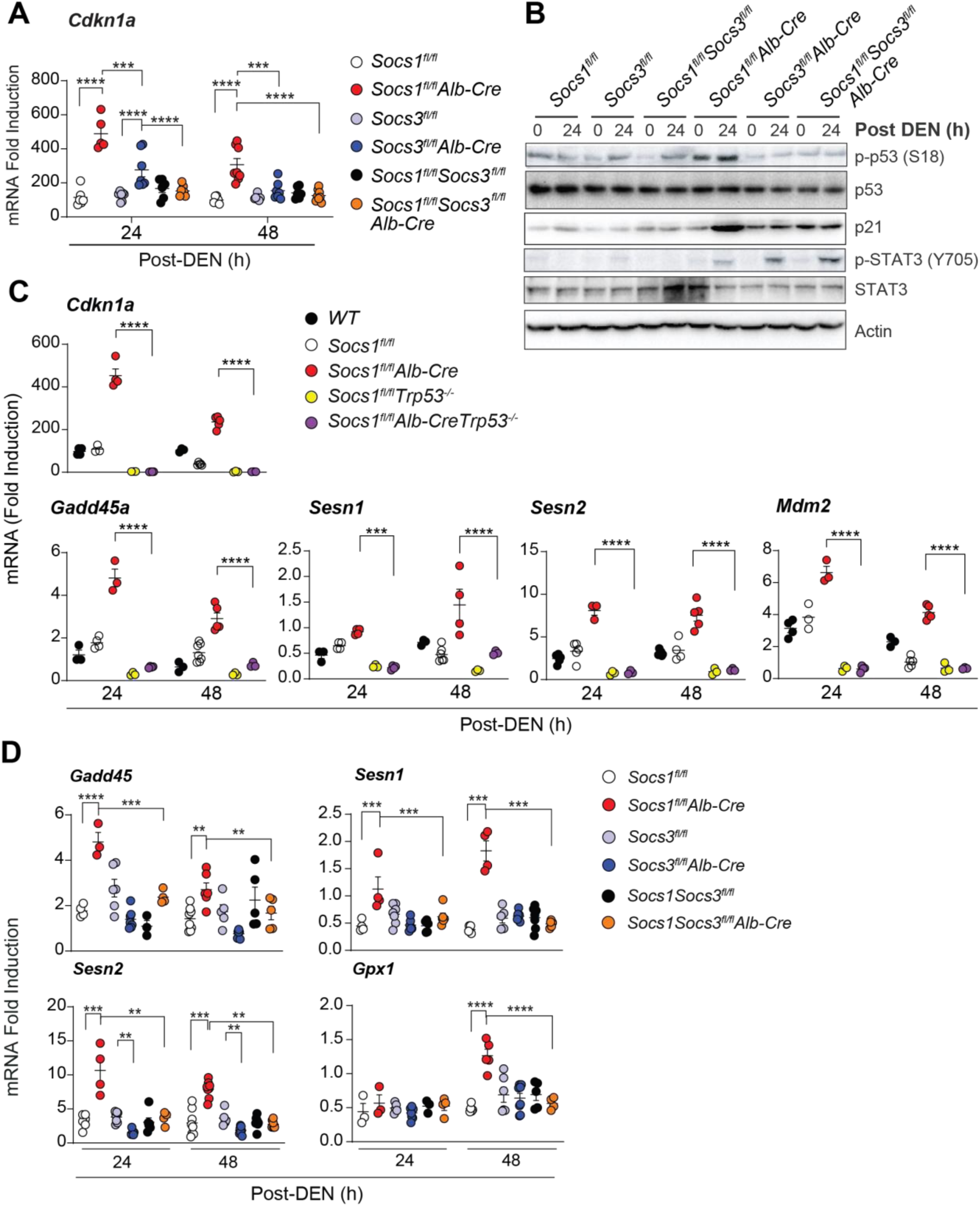
SOCS3 promotes p53-dependent induction of CDKN1A in SOCS1-deficient hepatocytes. (A) Hepatic *Cdkn1a* expression in mice lacking SOCS1, SOCS3 or both following DEN treatment (100 mg/Kg bodyweight; n=5-8 mice/group). mRNA fold induction was calculated relative to the expression in untreated mice of the same genotype after normalization to the housekeeping gene *Rplp0*. (B) Immunoblot analysis of phospho-p53, total p53, p21, phospho-STAT3, total STAT3 and actin in control and DEN-treated mice livers. Representative data from at least three experiments are shown. (C) Induction of *Cdkn1a* and the indicated p53 target genes in the livers of SOCS1-deficient mice lacking p53 (n=3-7 mice/group). (D) Induction of p53 target genes in the livers of mice lacking SOCS1, SOCS3 or both following DEN treatment was evaluated by qRT-PCR at 24 and 48 h later (n=3-6 mice/group). Statistics: Mean ± SEM; Two-way ANOVA with Tukey’s test; ** *p*<0.01, *** *p*<0.001, **** *p*<0.001.

DEN induces oxidative stress, DNA damage and sustained p53 activation in the liver ^26, 27^. As both SOCS1 and SOCS3 can activate p53 ^4–6^, we examined DEN-induced p53 activation in the livers of mice lacking SOCS1, SOCS3 or both. DEN-treated *Socs1^fl/fl^Alb-Cre* mice displayed increased p53 phosphorylation and CDKN1A (p21) protein expression in the liver, which was abrogated in *Socs1^fl/fl^Socs3^fl/fl^Alb-Cre* mice (**Figure 3B**). In contrast, STAT3 phosphorylation, presumably resulting from IL-6 signaling, was augmented by the loss of SOCS1, SOCS3 or both (**Figure 3B**). To determine whether the increased *Cdkn1a* induction in SOCS1-deficient livers required p53, we generated *Socs1^fl/fl^Alb-CreTp53^-/-^* mice and evaluated DEN-induced *Cdkn1a* expression. *Cdkn1a* induction in *Socs1^fl/fl^Alb-Cre* mice was abrogated by p53 deficiency (**Figure 3C**). Induction of other p53 target genes such as *Gadd45a, Sesn1, Sesn2* and *Mdm2*, which were also strongly upregulated in SOCS1-deficient livers, was also abolished by p53 loss (**Figure 3C**). Induction of these p53- dependent genes was compromised in *Socs1^fl/fl^Socs3^fl/fl^Alb-Cre* mice, similarly to *Cdkn1a* (**Figure 3A, 3D**). These data indicated that SOCS3 promoted p53 activation and CDKN1A upregulation in SOCS1-deficient hepatocytes.

### SOCS1 deficiency promotes SOCS3-dependent NFR2 activation

Neoplastic growth is often associated with increased cellular metabolism and oxidative stress ^28^. Cancer cells adapt to oxidative stress by upregulating NRF2, a transcriptional activator that induces antioxidant response genes including its own (*Nfe2l2*) ^21^. In normal cells, NRF2 is downregulated by Kelch-like ECH-associated protein 1 (KEAP1)-mediated ubiquitination and proteasomal degradation ^21^. The KEAP1-NRF2 interaction is disrupted by KEAP1 oxidation as well as by proteins such as CDKN1A and p62, which interact with NRF2 and KEAP1, respectively, leading to NRF2 activation ^19, 29^. To determine whether increased CDKN1A expression in SOCS1-deficient liver promoted NRF2 activation and whether this was mediated by SOCS3, we treated mice lacking SOCS1, SOCS3 or both with DEN, which induces oxidative stress in hepatocytes ^30^. DEN treatment markedly increased NRF2 gene (*Nfe2l2*) and protein expression in SOCS1-deficient livers that coincided with elevated p21 expression, both of which were abrogated by simultaneous loss of SOCS3 (**Figure 4A, 4B**). The increased NRF2 expression in SOCS1-deficient livers was associated with the induction of NRF2 target genes such as *Gstm1, Gstm4, Gclc, Gpx2, Hmox1* and *Nqo1*, whereas the expression of *Keap1* was not altered (**Figure 4A**). Next, we examined NRF2 expression in DEN-induced HCC nodules in mice lacking SOCS1, SOCS3 or both. Immunofluorescence staining of HCC nodules in *Socs1^fl/fl^Alb-Cre* mice showed elevated NRF2 expression compared to tumor nodules in control or SOCS3-deficient mice and this upregulation was abrogated in *Socs1^fl/fl^Socs3^fl/fl^Alb-Cre* mice (**Figure 4C, 4D**; **Figure S3**). HCC nodules of SOCS1-deficient mice also showed increased expression of *Cdkn1a*, *Nfe2l2* and the NRF2 target genes *Gstm4, Gclc* and *Nqo1*, all of which were attenuated in tumors from *Socs1^fl/fl^Socs3^fl/fl^Alb-Cre* mice (**Figure 4E**). These findings indicated that SOCS3 upregulates NRF2 expression and its transcriptional activity in SOCS1-deficient hepatocytes under conditions of increased oxidative stress.

**Figure 4.**
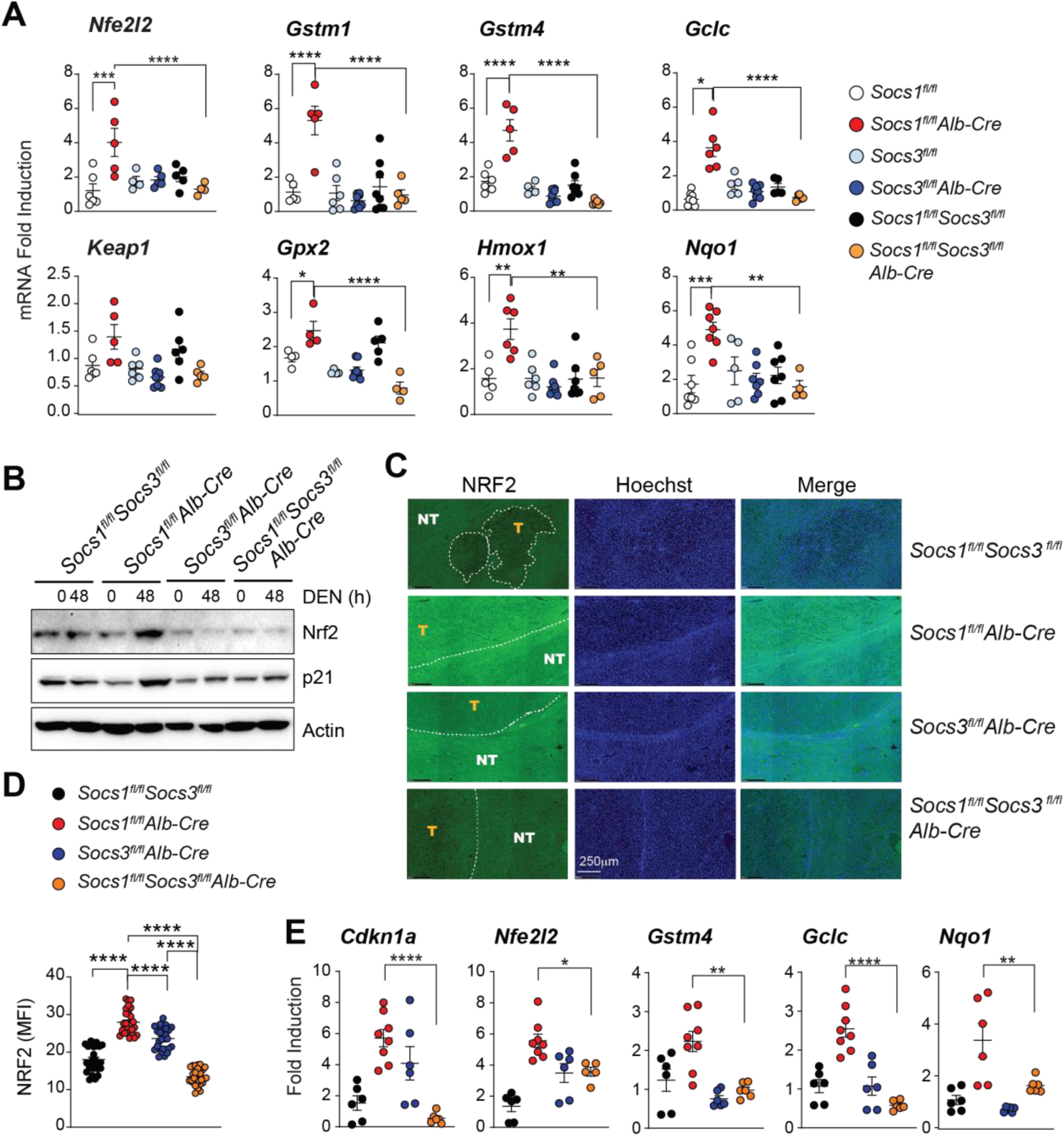
SOCS1 deficiency upregulates NRF2 target genes in a SOCS3-dependent manner. (A) Induction of *Nfe2l2* and NRF2 target genes in mice lacking SOCS1, SOCS3 or both in hepatocytes 48h after DEN treatment (n=4-8 mice/group). (B) NRF2 and p21 protein expression in the liver tissues of DEN-treated mice lacking SOCS1, SOCS3 or both in hepatocytes. Representative data from 3 mice/ genotype/ group. (C) NRF2 immunostaining in the tumor bearing livers. Tumor nodules (T) and surrounding normal tissue (NT) are demarcated (10X magnification). Representative images from 3 mice per group. (D) Mean fluorescence intensity of NRF2 staining within the tumor nodule (n=3 mice/group) (E) Expression of *Cdkn1a, Nfe2l2* and NRF2 target genes in microscopically dissected HCC tumor nodules (T) compared among different genotypes. (n=6-8 mice/group). Statistics: Mean ± SEM; One-way ANOVA with Tukey’s multiple comparison test; \**p*<0.05, \*\**p*<0.01, \*\*\**p*<0.001, \*\*\*\**p*<0.0001.

### NFR2 activation in SOCS1-deficient hepatocytes requires CDKN1A

To determine whether SOCS3-dependent NRF2 activation in SOCS1-deficient hepatocytes requires CDKN1A, we generated *Cdkn1a^-/-^* mice lacking SOCS1 in hepatocytes. DEN induced strong NRF2 immunostaining in control mice livers that was markedly enhanced by SOCS1 deficiency (**Figure 5A**, ii versus iv) and this increase was profoundly attenuated by simultaneous loss of CDKN1A (**Figure 5A**, iv versus vi). The CDKN1A-dependent increase in NRF2 expression in SOCS1-deficient livers was also confirmed by western blot (**Figure 5B**). *Nfe2l2* and the NRF2 target genes were strongly induced in *Socs1^fl/fl^Alb-Cre* mice, whereas this induction was significantly diminished in *Socs1^fl/fl^Alb- CreCdkn1a^-/-^* mice (**Figure 5C**). These findings demonstrate that activation of the SOCS3-p53- CDKN1A axis in SOCS1-deficient hepatocytes leads to increases NRF2 expression and its transcriptional activity.

**Figure 5.**
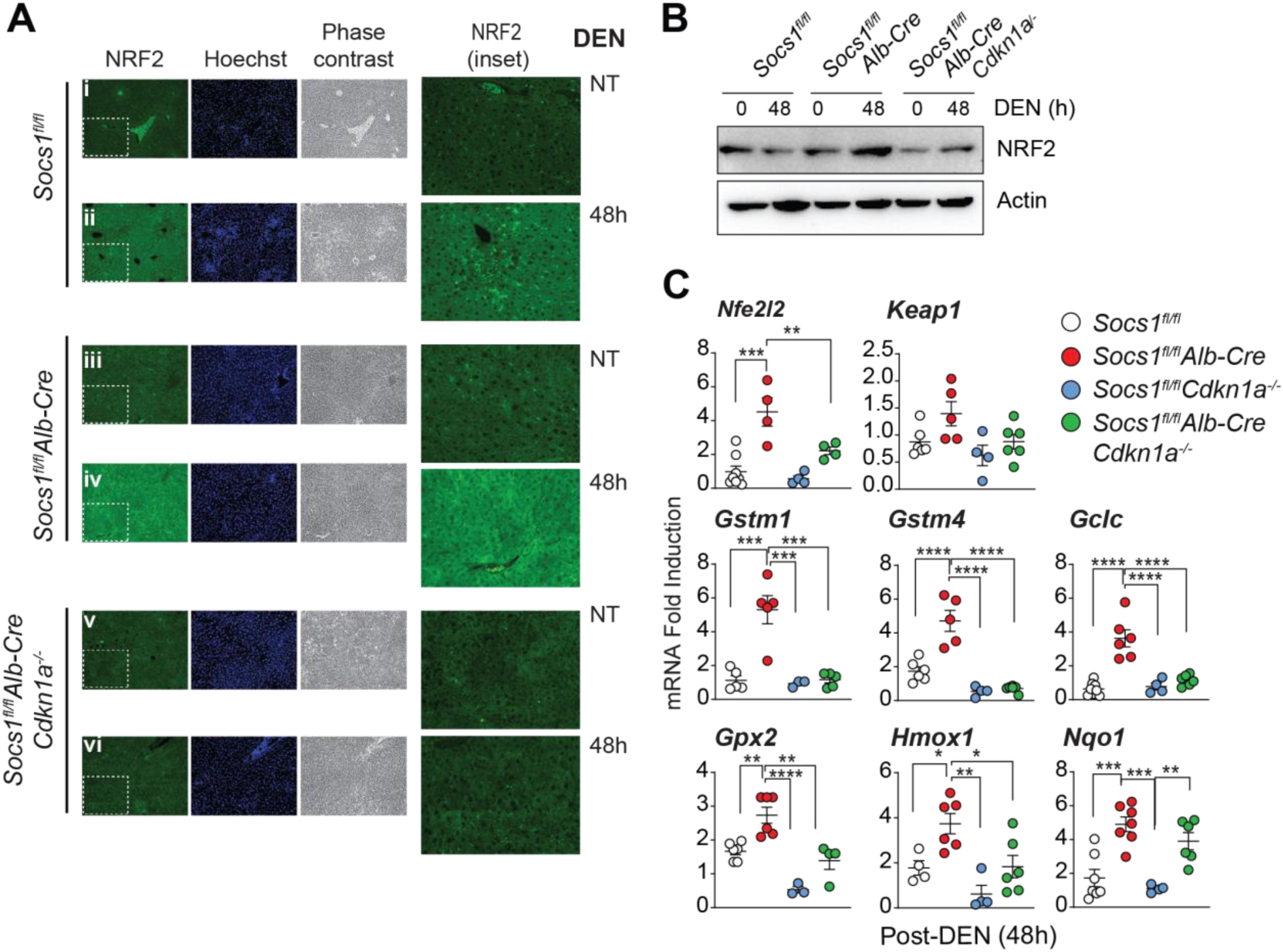
CDKN1A drives the induction of NRF2 target genes and increases the ability to tolerate oxidative stress in SOCS1-deficient livers. (A) NRF2 expression in the livers of the indicated mice groups after DEN treatment. 10X magnification. NRF2 staining in the inset is digitally magnified in the right column. (B) NRF2 protein expression in liver lysates. Representative data from 3 mice/group are shown for A and B. (C) *Nfe2l2* and NRF2 target gene expression in the livers 48h after DEN treatment. Cumulative data from 4-8 mice/group. Gene expression data for *Socs1^fl/fl^* and *Socs1^fl/fl^Alb-Cre* are duplicated from Figure 5F-G for comparison. Statistics: Mean ± SEM; One-way ANOVA with Tukey’s multiple comparison test; \**p*<0.05, \*\**p*<0.01, \*\*\**p*<0.001, \*\*\*\**p*<0.0001.

### CDKN1A drives carcinogenesis in SOCS1-deficient hepatocytes

Increased oxidative stress leads to lipid peroxidation, which generates 4-hydroxy-2-nonenal (4-HNE), that amplifies the oxidative stress ^31, 32^. Liver sections from DEN-treated wildtype mice displayed increased 4-HNE staining, which was significantly increased by SOCS1 deficiency, and this increase was attenuated by simultaneous ablation of *Cdkn1a* (**Figure 6A, 6B**; **Figure S4A**). These data indicated that CDKN1A enables SOCS1-deficient hepatocytes to withstand increased oxidative stress. Next, we evaluated DEN-induced HCC in SOCS1-deficient livers lacking *Cdkn1a*. CDKN1A-deficient mice showed an increased HCC incidence and tumor volume compared to control mice (**Figure 6C, 6D**; **Figure S4B**), which supported the tumor suppressor function of CDKN1A. However, *Socs1^fl/fl^Alb-Cre Cdkn1a^-/-^* mice showed reduced HCC incidence with significantly fewer and smaller tumor nodules compared to *Socs1^fl/fl^Alb-Cre* mice (**Figure 6C, 6D**; **Figure S4B**). These results demonstrated that CDKN1A is an obligate mediator of increased tolerance to oxidative stress and enhanced hepatocarcinogenesis in SOCS1-deficient livers.

**Figure 6.**
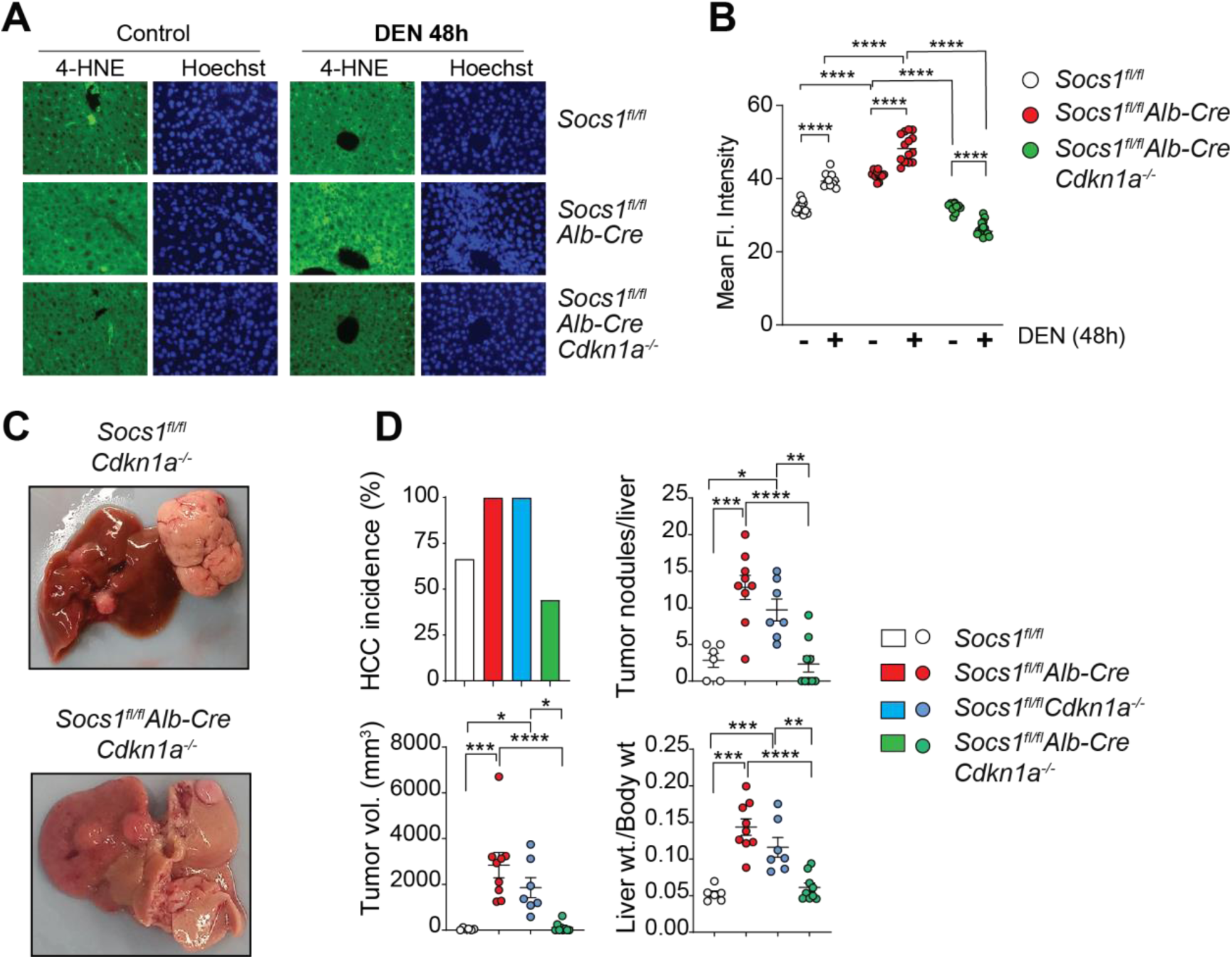
CDKN1A is necessary for increased lipid peroxidation and HCC development in SOCS1-deficient livers. (A) 4-HNE staining for lipid peroxidation in DEN-treated mice livers. Representative images from 3 mice/group are shown at 40X magnification. (B) Quantification of 4-HNE staining (n=3 mice/group). (C,D) HCC development in SOCS1-deficient mice lacking CDKN1A and control CDKN1A-deficient mice 10 months after treatment with DEN (25 mg/Kg bodyweight) at 2 weeks of age. Macroscopic liver images, representative of n=6-9 mice per group are shown. (D) Cumulative data on HCC incidence, number of visible tumors, tumor volume (sum of all tumors ≥2 mm in diameter) and liver to bodyweight ratio. For comparison, HCC data for *Socs1^fl/fl^* and *Socs1^fl/fl^Alb-Cre* mice are duplicated from Fig 2C. Statistics: Mean ± SEM; One-way (D) or two-way (B) ANOVA with Tukey’s multiple comparison test: \**p*<0.05, \*\**p*<0.01, \*\*\**p*<0.001, \*\*\*\**p*<0.0001. n.s., not significant.

### Elevated *CDKN1A* and NRF2 target gene expression in *SOCS1-low/SOCS3-high* HCC

Next, we determined how *SOCS1* and *SOCS3* expression correlated with CDKN1A expression in human HCC. Both *SOCS1-high* and *SOCS3-high* groups showed elevated *CDKN1A* expression (**Figure 7A**) and *SOCS1-low*/*SOCS3-high* HCC cases showed significantly elevated *CDKN1A* expression compared to the *SOCS1-low/SOCS3-low* group (**Figure 7B**). We performed a gene set enrichment analysis (GSEA) for gene ontology (GO) terms containing ‘oxygen’ and ‘oxidant’ for each of the HCC groups compared to normal liver tissues (**Figure 7C**). The *SOCS1-low/SOCS3-low* group showed significant negative enrichment for antioxidant activity genes with a normalized enrichment score (NES) of -1.6588757 (*p*=3e-04; *p*.*adjusted*=7e-04) whereas the *SOCS1-low/SOCS3- high* group showed a positive enrichment for cellular response to increased oxygen levels (NES=1.7385050; *p*=1e-04; *p*.*adjusted*=0.0011) (**Figure 7D**). Importantly, the benchmark NRF2 target gene signature curated by Polonen and colleagues ^33^ revealed significant enrichment in *SOCS1- low/SOCS3-high* (NES=1.4661141; *p*=1e-04; *p*.*adjusted*=5e-04) and *SOCS1-high/SOCS3-high* (NES=1.471197; *p*=0.021; *p*.*adjusted*=0.0104) groups (**Figure 7D, 7E**). The *SOCS1-high/SOCS3-low* group, however, did not show any enrichment for antioxidant activity genes (**Figure S5**). These data indicated that elevated expression of SOCS3 and CDKN1A in SOCS1-deficient HCC could confer protection against oxidative stress and thus contribute to aggressive cancer.

**Figure 7.**
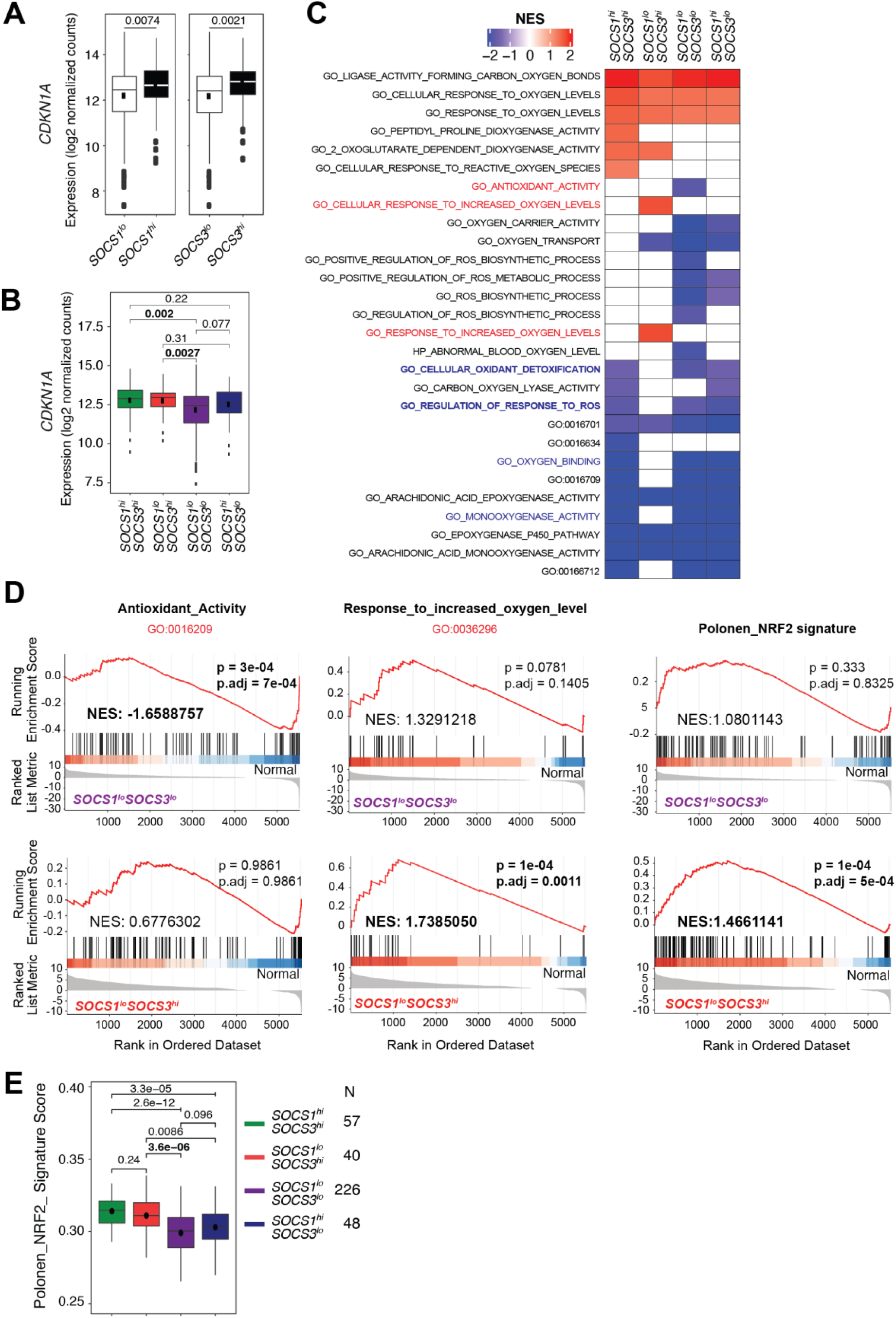
*SOCS1-low/SOCS3-high* HCC cases display elevated CDKN1A expression and enrichment of NRF2 signature genes. (A) *CDKN1A* expression in TCGA-LIHC cases dichotomized based on *SOCS1* or *SOCS3* mRNA levels (z score). (B) Increased *CDKN1A* expression in *SOCS1-low/SOCS3-high* HCC than in *SOCS1-low/SOCS3-low* group. (C) Gene set enrichment analysis (GSEA) for biological pathways containing the term ‘oxygen’ and ‘oxidant’ in the indicated HCC groups compared to normal liver tissues. The description of GO terms identified by number are indicated in supplementary Fig. S4. (D) GSEA plots showing negative enrichment of Antioxidant_Activity’ pathway genes in *SOCS1- low/SOCS3-low* group and positive enrichment of ‘Response_to_increased_oxygen_level’ pathway and ‘Polonen_NRF2 Signature’ genes in the *SOCS1-low/SOCS3-high* group. (E) Increased enrichment of the ‘Polonen_NRF2 Signature’ genes in the *SOCS1-low/SOCS3-high* group than in the *SOCS1-low/SOCS3-low* group.

### SOCS1 limits the ability to tolerate oxidative stress by inhibiting NRF2

We have previously shown that SOCS1 expression in HCC cell lines reduces CDKN1A expression and *in vivo* tumor growth ^10, 15^. To test the impact of SOCS1 on cellular antioxidant response, we treated murine Hepa1-6 (Hepa) hepatoma cells expressing SOCS1 (Hepa-SOCS1) or empty vector (Hepa-vector) with *tert*-butyl hydroperoxide (*t*-BHP) or cisplatin, which induce oxidative stress ^34, 35^. Hepa-SOCS1 cells treated with *t*-BHP or cisplatin showed reduced survival compared to Hepa-vector cells (**Figure 8A**; **Figure S6A**). Fluorescence indicators CellROX Green and CellROX Deep Red, which detect mitochondrial/nuclear and cytoplasmic ROS respectively, showed markedly reduced ROS levels in Hepa-SOCS1 cells compared to Hepa-vector cells following *t*-BHP treatment (**Figure 8B**). Immunostaining of 4-HNE was also markedly reduced in Hepa-SOCS1 cells (**Figure 8C**).

**Figure 8.**
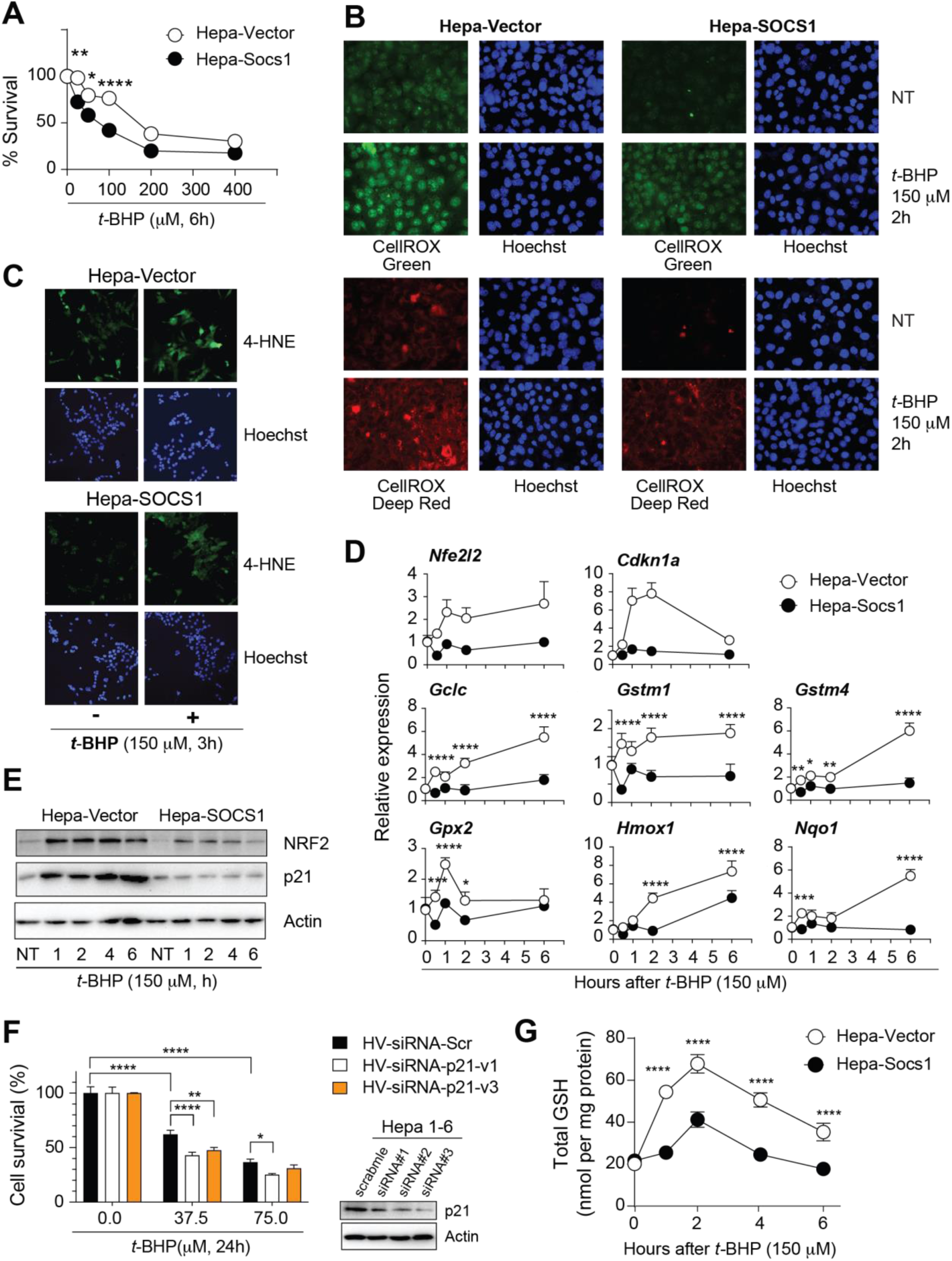
SOCS1 limits the ability to tolerate oxidative stress by inhibiting NRF2. (A) Decreased survival of Hepa 1-6 cells expressing SOCS1 (Hepa-SOCS1) compared to control cells (Hepa-vector) 6 h after treatment with the indicated concentrations of *t*-BHP. Representative data from more than three experiments done in triplicates are shown. (B) Representative images of mitochondrial/nuclear and cytosolic ROS levels (40X magnification) in Hepa-vector and Hepa-SOCS1 cells treated with *t*-BHP. (C) Representative 4-HNE staining of lipid peroxidation (10X magnification) in Hepa-vector and Hepa-SOCS1 cells treated with *t*-BHP. Representative images from at least three experiments are shown for B and C. (D, E) Induction of *Nfe2l2*, *Cdkn1a* and NRF2 target genes (n=4) in Hepa-vector and Hepa-SOCS1 cells treated with *t*-BHP for the indicated periods. (E) Protein levels of NRF2 and p21 in *t*-BHP-treated Hepa-vector and Hepa-SOCS1 cells. (F) Survival of Hepa-Vector cells transfected with *Cdkn1a*-targeting siRNA 24h prior to *t*-BHP treatment (n=3). (G) Glutathione levels in *t*-BHP-treated Hepa-vector and Hepa-SOCS1 cells (n=3). Statistics: Mean ± SEM; Two-way ANOVA with Tukey’s multiple comparison test; \**p*<0.05, \*\**p*<0.01, \*\*\**p*<0.001, \*\*\*\**p*<0.0001.

As increased ROS levels are usually associated with cellular damage, the reduced survival of Hepa- SOCS1 cells despite lower ROS levels and lipid peroxidation (**Figure 8A-C**) raised a conundrum, which could result from the destabilizing effect of SOCS1 on CDKN1A and the consequent reduction in the ability to activate NRF2. Following exposure to *t*-BHP, Hepa-vector cells displayed strong induction of *Nfe2l2* gene and protein expression within 1h that remained high up to 6h, whereas these events were markedly diminished in Hepa-SOCS1 cells (**Figure 8D,E**). This was accompanied by a concomitant reduction in CDKN1A mRNA and protein levels in Hepa-SOCS1 cells (**Figure 8D,E**). Similar changes in *Nfe2l2* and *Cdkn1a* gene and protein expression were observed following cisplatin treatment of Hepa-vector and Hepa-SOCS1 cells (**Figure S6B,C**). siRNA-mediated knockdown of *Cdkn1a* in Hepa-vector cells significantly reduced cell survival following exposure to *t*-BHP (**Figure 8F**), indicating an important role for CDKN1A in promoting cell survival despite high levels of ROS (**Figure 8B**). Human Hep3B cells expressing SOCS1 also showed reduced survival and expression of NRF2 and p21 (**Figure S6D,E**). Hepa-SOCS1 cells exposed to *t*-BHP or cisplatin showed significantly lower induction of NRF2-induced antioxidant response genes *Gclc, Gstm1, Gstm4, Gpx2, Hmox1* and *Nqo1* (**Figure 8D**; **Figure S6F**). Accordingly, *t*-BHP markedly increased glutathione levels in Hepa-vector cells, whereas this adaptation was significantly reduced in Hepa-SOCS1 cells (**Figure 8G**). These results indicate that SOCS1 inhibits the ability of CDKN1A to activate NRF2-mediated antioxidant response that sustains increased cellular metabolism and further ROS production in HCC.

## Discussion

In this study we show that even though SOCS1, SOCS3 and CDKN1A individually function as tumor suppressors in hepatocytes, SOCS3 and CDKN1A gain oncogenic potential in SOCS1- deficient hepatocytes via promoting NRF2-mediated cellular adaptation to oxidative stress leading to HCC progression (**Figure 9**). Our findings also suggest that targeting the NRF2-mediated tumorigenic pathway could be a potential drug target to treat *SOCS1-low* HCC.

**Figure 9.**
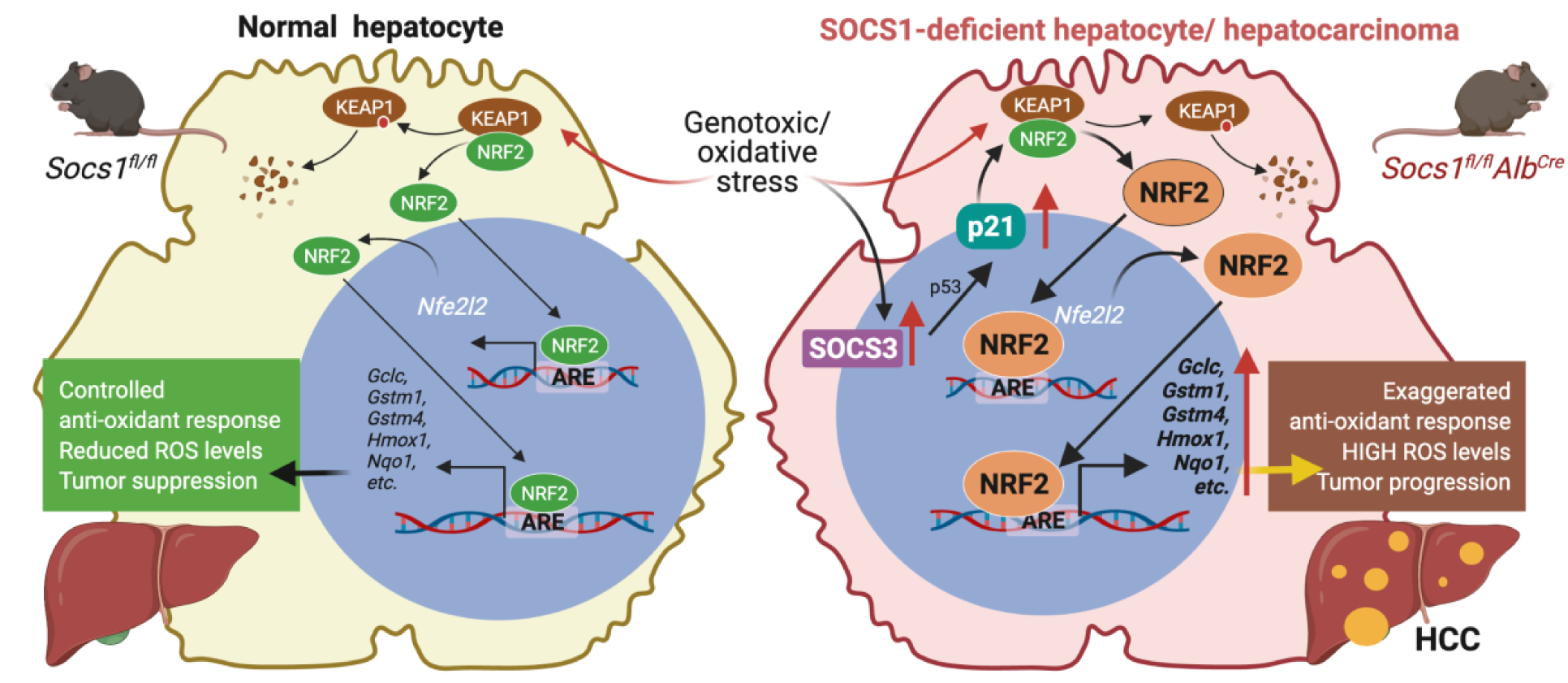
Proposed molecular mechanism underlying increased HCC development in SOCS1- deficient livers. In SOCS1-deficient hepatocytes, genotoxic stress upregulates SOCS3, which activates p53 that induces CDKN1A expression. CDKN1A in turn promotes transcriptional activation of NRF2, expression of NRF2 target genes and cellular adaptation to increased oxidative stress to support neoplastic growth.

Clinical data and genetic models support independent tumor suppressor functions of SOCS1 and SOCS3 in the liver ^7–9, 11, 12^. SOCS3 is crucial to control IL-6 signaling, whereas SOCS1 regulates HGF signaling in the liver ^12–15^. Both SOCS1 and SOCS3 promote transcriptional activation of p53 ^4–6^, suggesting that loss of SOCS1 or SOCS3 could compromise the tumor suppressor functions of p53. However, in the liver, SOCS1 deficiency did not attenuate p53 activation in response to DEN ^10^, whereas the combined loss of SOCS1 and SOCS3 compromised the induction of p53 target genes. The current investigation revealed that regulating the tumor promoting CDKN1A-mediated NRF2 activation is a crucial tumor suppression function of SOCS1 in HCC.

CDKN1A generally functions as a tumor suppressor by inhibiting CDK1 and CDK2 thereby blocking cell cycle progression, and CDKN1A-deficiency promotes spontaneous and induced tumors in different tissues ^17^. Ablation of *Cdkn1a* in hepatocyte-specific NEMO-deficient mice, which develop chronic hepatitis, leads to spontaneous HCC ^36^. We also show in our study that CDKN1A deficiency increases DEN-induced HCC. However, elevated CDKN1A expression occurs in many cancers including HCC and correlates with poor prognosis, suggesting an oncogenic role of CDKN1A ^17, 37^. Indeed, Vogel and colleagues have shown that CDKN1A is required for HCC development under conditions of mild inflammation in *Fah^-/-^* mice and following chronic cholestatic liver injury in *Mdr2^-/-^* mice ^37, 38^. These findings showed that CDKN1A is an ‘oncojanus’ that can either inhibit or promote HCC depending on the molecular context. CDKN1A gains oncogenic potential when retained in the cytosol via AKT-dependent phosphorylation where it interferes with apoptotic mediators ^17^. CDKN1A can also promote CDK4 activity and facilitate G1 phase progression ^39^. CDKN1A promotes HCC in *Mdr2^-/-^* mice likely via facilitating CyclinD-CDK4 complex formation and cell cycle progression ^37^. Even though the oncojanus role of CDKN1A is firmly established, conditions that promote its oncogenic potential are poorly understood. Epigenetic repression of *SOCS1* in HCC can contribute to CDKN1A-mediated oncogenesis by at least two mechanisms. Increased AKT activation downstream of deregulated growth factor signaling in SOCS1-deficient livers results in CDKN1A phosphorylation and cytosolic retention ^10^. As shown in the present study, compensatory increase in SOCS3 expression can lead to increased *CDKN1A* expression via p53 activation. Even though disabling p53 mutations occur in a third of non-aflatoxin-induced HCC ^40^, the oncogenic pathway driven by the SOCS3-p53-CDKN1A axis could still occur in a considerable proportion of HCC cases with *SOCS1* repression but with intact *SOCS3* gene. Supporting this notion, *SOCS1-low/SOCS3-high* cases in the TCGA-LIHC cohort show elevated *CDKN1A* expression and poor progression-free survival.

Cytosolic CDKN1A can bind NRF2 and relieve the inhibitory KEAP1-NRF2 interaction leading to its activation ^19, 21^. NRF2 is well known as an oncojanus as it enables cancer cells to cope with increased oxidative stress during cancer progression ^21^. NRF2 is also activated by the selective autophagy substrate p62 (SQSTM1), which disrupts NRF2-KEAP1 interaction ^29^. Reduced HCC development in *Fah^-/-^* mice was attributed to compensatory induction of Sestrin2, which promoted p62-dependent autophagic degradation of KEAP1, leading to NRF2 activation, protection from oxidative damage and neoplastic transformation ^38^. Contrary to the findings in *Cdkn1a^-/-^Fah^-/-^* mice, *Cdkn1a^-/-^Socs1^-/-^* mice showed impaired induction of NRF2 target genes. While CDKN1A attenuates the antitumor functions of NRF2 in FAH-deficient livers ^38^, our findings indicate that CDKN1A promotes the oncogenic potential of NRF2 in SOCS1-deficient livers. These contrasting effects of CDKN1A-mediated NRF2 activation likely reflect the duality of NRF2 functions, being protective in normal and preneoplastic stages but detrimental in transformed hepatocytes ^41^.

NRF2-deficient mice have clearly established an oncogenic role for NRF2 activation in DEN-induced HCC ^42^ and NRF2-high HCC patients show poor survival ^43, 44^. Even though mutations that disrupt KEAP-NRF2 interactions contribute to NRF2 activation in human and rodent HCC ^45, 46^, NRF2 activation in HCC can also occur by other mechanisms such as elevated expression of p62 ^47^. Our findings add to this list CDKN1A induced by SOCS1 deficiency. Besides conferring protection from oxidative stress, NRF2 activation in hepatocytes can also provide growth stimuli via AKT activation and induction of PDGF and EGF that could promote carcinogenesis ^48^. As growth factor-induced AKT activation contributes to CDKN1A phosphorylation and its cytosolic retention ^10^, NRF2 can establish a positive feedback loop in CDKN1A-overexpressing cancers. SOCS1 deficiency can doubly contribute to this amplification loop by deregulating growth factor signaling as well as upregulating CDKN1A and NRF2.

Our findings unravel the complexities of targeting individual genetic aberrations or signaling pathways in cancers that could explain the limited achievements of targeted therapeutics in HCC ^23^. Our findings also highlight the potential usefulness of considering genetic defects and molecular pathway signatures together to predict and test the context-dependent alterations in protein functions that could pave way for a more effective precision therapeutics. Even though we have revealed the pro-tumorigenic potential of CDKN1A and NRF2 in the context of SOCS1deficient HCC, targeting these aberrant functions without compromising their antitumor activities, or restoring SOCS1 will be challenging. To overcome these limitations, anticancer drugs activated by NRF2-induced enzymes such as NQO1 are being developed ^49^, which could be exploited to treat *SOCS1-low* HCC with high antioxidant response gene signature in the foreseeable future.

## Materials and methods

### TCGA-LIHC dataset and analyses

Transcriptomic data on The Cancer Gene Atlas-liver HCC (TCGA-LIHC) study cohort and the associated clinicopathological information ^24^ were downloaded (http://tcga-data.nci.nih.gov/tcga/) using Bioconductor package TCGAbiolinks_2.14.1 ^50^. TCGA level 3 data comprised of 50 normal tissue, 371 primary tumors were used after excluding three recurrent tumors.

#### Gene expression analysis

RNAseq read counts downloaded from TCGA-LIHC were normalized for sequencing depth using the size factor method implemented in Deseq2_1.26.0 package ^51^. Log2 normalized read counts were used to show gene expression levels. Significant difference in gene expression between groups was measured by Wilcoxon test at *p* <0.05.

#### Survival analysis

To conduct survival analysis, *SOCS1* and *SOCS3* expression were converted to z-score and used to divide patients into high expression and low expression groups. Patients were further stratified into four groups combing *SOCS1* and *SOC3* expression. Kaplan-Meir survival plots were generated using the R packages *survival_3.1-12* and *survminer_0.4.6* ^52, 53^. Disease-free survival was compared between the four groups using log-rank test. The hazard ratio was calculated via Cox regression model using *survival_3.1-12* ^52^.

#### Pathway analysis

“Oxygen” and “oxidant” related gene set in the gene ontology Biological process (GO: BP) were downloaded from the Molecular Signatures database (MsigDB) ^54^ using *msigdbr_7.2.1* package ^55^. A mod.t.test function (MKmisc 1.6 R package) ^56^ was used to compare each of the four patient groups (segregated based on the expression levels of *SOCS1* and *SOCS3*) to benign samples and to score genes. Genes were then rank ordered and gene enrichment analysis was performed using *clusterProfiler_3.14.3* R package ^57^. Gene sets were considered enriched with a Benjamini-Hochberg (BH) adjustment <0.05. Enrichment of NRF2 gene signature benchmarked by Polonen and colleagues (Ref. 26) was analyzed using the singscore 1.6.0_R package ^58^.

#### Statistical analysis

All data analysis and statistical tests were performed in R version 3.6.2 (2019- 12-12).

### Mouse strains

Hepatocyte-specific SOCS1-deficient mice (*Socs1^fl/fl^Alb^Cre^*) were previously described ^10^. *Socs3^fl/fl^*, *Alb^Cre^*, *Cdkn1a^-/-^* and *p53^-/-^* mice were purchased from the Jackson Laboratory. Mice lacking SOCS3 or both SOCS1 and SOCS3 in hepatocytes (*Socs3^fl/f^Alb^Cre^*; *Socs1^fl/fl^Socs3^fl/fl^Alb^Cre^*) were generated for this study. *Socs1^fl/fl^Alb^Cre^* mice were crossed with *Cdkn1a^-/-^* or *Tp53^-/-^* to generate SOCS1- deficient mice also lacking CDKN1A (*Socs1^fl/fl^Alb^Cre^Cdkn1a^-/-^*) or p53 (*Socs1^fl/fl^Alb^Cre^Tp53^-/-^*) in hepatocytes. All mice strains used in this study are in C57BL/6N background and are listed in **Supplementary Table S1**. Control mice were derived from littermates. Mice were housed in ventilated cages with 12 hours day/night cycle and fed with normal chow *ad libitum*. All experiments on mice were carried out during daytime with the approval of the Université de Sherbrooke Ethics Committee for Animal Care and Use (Protocol ID 226-17B; 2017-2043).

### Experimental HCC

To induce HCC, DEN was administered via intraperitoneal (i.p.) route (25mg/kg bodyweight) into two weeks-old male mice as females are resistant to DEN-induced HCC ^10, 59^. All reagents and their sources are listed in **Supplementary Table S2**. Treated mice were sacrificed 10 months later and visible tumor nodules were counted. Tumor dimensions were measured using a digital Vernier caliper. Tumor volume was calculated in mm^3^ using the formula: (length x width^2^)/2. Liver tissues were collected in buffered formalin for histology.

### Induction of genotoxic and oxidative stress

To induce oxidative and genotoxic stress in the liver, 6-8 weeks-old male mice were injected DEN (100mg/kg bodyweight, i.p.) ^10^. At the indicated time points, liver tissues were fixed in buffered formalin, preserved in RNAlater® for gene expression analysis or snap-frozen and stored at -80°C to evaluate protein expression.

### Immunofluorescence staining of liver sections

To assess cell proliferation within tumor nodules, liver sections were immunostained for Ki67 and immunofluorescence (IF) images captured by NanoZoomer and analyzed by NanoZoomer Digital Pathology (NDP) view2 software (Hamamatzu Photonics, Japan). Software source and versions are listed in **Supplementary Table S3** and antibodies used for IF in **Supplementary Table S4**. The number of Ki67+ nuclei were counted in 8-10 random fields for each specimen. NRF2 expression in tumors was evaluated by IF and mean fluorescence intensity (MFI) was quantified (8-10 random fields/section; 3-5 mice per genotype) using the Image J software (National Institutes of Health, USA). Lipid peroxidation in DEN-treated liver tissues was evaluated by IF staining of 4- hydroxynonenol (4-HNE). Images were captured using the Axioskop 2 microscope (Carl Zeiss, Inc.) and MFI was quantified as for NRF2 using pooled data from 3-5 mice per group.

### Cell culture and assays

Hepa1-6 (Hepa) murine hepatoma and Hep3B human HCC cell lines were purchased from ATCC. Stable lines of Hepa and Hep3B cells expressing SOCS1 (Hepa-SOCS1) or the control vector were previously described ^14, 15^. Details of the cell lines are given in **Supplementary Table S5**. Hepa cells were grown in Dulbecco’s modified Eagle’s medium (DMEM) containing 10% fetal bovine serum (FBS). Hep3B cells were cultured in DMEM containing 10% FBS. To induce oxidative stress, Hepa and Hep3B cells were treated with the indicated concentrations of *tert*-butyl hydroperoxide (*t*-BHP) or cisplatin (*cis*-diamino-dichloroplatinum). Cell survival was measured using the WST-8 (water-soluble Tetrazolium-8: 2-(2-methoxy-4-nitrophenyl)-3-(4-nitrophenyl)- 5- (2,4-disulfophenyl)-*2H*-tetrazolium) assay kit (CCK-8; Dojindo Molecular Technologies, #CK04) following manufacturer’s instructions. Briefly, 2500-5000 cells were seeded in 96-well plates in 100μl medium in the presence or absence of *t*-BHP or *cis*-Pt. After the indicated periods of incubation, 100μl WST-8 reagent was added to each well and incubated for 2 hours. Color development, indicative of cell number and viability, was measured at 440nm wavelength using SPECTROstar^Nano^ (BMG Labtech, Germany) spectrometer. Oxidative stress was evaluated using the CellROX Green and CellROX Deep Red fluorescent ROS indicators (ThermoFisher). Silencing of *Cdkn1a* in Hepa cells was achieved by siRNA (**Supplementary Table S6**) transfection using with Lipofectamine RNAiMAX from Invitrogen. Total glutathione (GSH) content was measured using the Glutathione Colorimetric Detection Kit (Invitrogen) following manufacturer’s instructions.

### Gene and protein expression analysis

Total RNA was isolated using RNeasy® kit from cultured hepatocytes and livers fixed in RNAlater®. Real-time quantitative PCR analysis was done as described (Ref. 4) using primers listed in **Supplementary Table S7**. Gene expression in each DEN-treated mouse was normalized to the reference gene *Rplp0* and mRNA fold-induction was calculated relative to the expression in untreated mice of the same genotype. Preparation of cell and liver tissue lysates and western blot have been previously described (Ref. 4). Antibodies used for western blotting are listed in **Table S4**.

### Statistical Analysis

Data were analyzed using the GraphPad Prism (San Diego, CA, USA) and represented as mean ± standard error of mean (SEM). Statistical significance was calculated by one-way or two-way ANOVA with Tukey’s multiple comparison test, and *p* values are represented by asterisks: * <0.05, ** <0.01, *** <0.001, **** <0.001.

## Abbreviations

*t*-BHP: *tert*-Butyl hydroperoxide
DEN: diethylnitrosamine
GO: gene ontology
4-HNE: 4-hydroxynonenal
NES: normalized enrichment score
ROS: reactive oxygen species

## Supplementary data

Supplementary Figures S1-S6 and Supplementary Tables S1-S7 are provided in the supplementary information associated with this manuscript.

## Acknowledgements

MGMK and MY were supported by graduate scholarships from FRQNT. AG received a postdoctoral fellowship and RK graduate scholarship from FRQS. RB is a recipient of the ‘Abdenour-Nabid, MD’ scholarship from Université de Sherbrooke. DPL is an FRQS-Junior-1 Research Scholar. CR-CHUS is an FRQS-funded research center.

## Author contributions

MGMK: Design and implementation of main experiments, Consolidation of figures, Manuscript writing- first draft; NB: TCGA-LIHC data analysis, Manuscript- review and editing; MY: Original data, Hypothesis generation; AG, RK and RB: Fluorescence microscopy and RT-qPCR analyses; SP: Analysis of histology data; AS: TCGA-LIHC data analysis, Manuscript- review and editing; AY: Generation of *Socs1*-floxed mice, Project development, Manuscript- review and editing; SR: generation of mouse strains, Data analysis and interpretation, Manuscript- review and editing; GF: Conceptualization, Data interpretation, Manuscript- review and editing; DPL: TCGA-LIHC data analysis, Manuscript- review and editing; SI: Conceptualization, Securing funding, Supervision, Data analysis and interpretation, Manuscript- writing, review and editing. All authors have read and approved the final manuscript.

## References

1. Yoshimura A, Ito M, Chikuma S, Akanuma T, Nakatsukasa H. Negative Regulation of Cytokine Signaling in Immunity. Cold Spring Harb Perspect Biol 10, (2018).

2. Kazi JU, Kabir NN, Flores-Morales A, Ronnstrand L. SOCS proteins in regulation of receptor tyrosine kinase signaling. Cellular and molecular life sciences : CMLS 71, 3297–3310 (2014).

3. Fujimoto M, Naka T. SOCS1, a Negative Regulator of Cytokine Signals and TLR Responses, in Human Liver Diseases. Gastroenterol Res Pract 2010, 470468 (2010).

4. Calabrese V, et al. SOCS1 links cytokine signaling to p53 and senescence. Mol Cell 36, 754–767 (2009).

5. Kong X, et al. Interleukin-22 induces hepatic stellate cell senescence and restricts liver fibrosis in mice. Hepatology 56, 1150–1159 (2012).

6. Saint-Germain E, et al. Phosphorylation of SOCS1 Inhibits the SOCS1-p53 Tumor Suppressor Axis. Cancer Res 79, 3306–3319 (2019).

7. Yoshikawa H, et al. SOCS-1, a negative regulator of the JAK/STAT pathway, is silenced by methylation in human hepatocellular carcinoma and shows growth- suppression activity. Nat Genet 28, 29–35. (2001).

8. Yang B, Guo M, Herman JG, Clark DP. Aberrant promoter methylation profiles of tumor suppressor genes in hepatocellular carcinoma. Am J Pathol 163, 1101–1107 (2003).

9. Yoshida T, et al. SOCS1 is a suppressor of liver fibrosis and hepatitis-induced carcinogenesis. J Exp Med 199, 1701–1707 (2004).

10. Yeganeh M, et al. Suppressor of cytokine signaling 1-dependent regulation of the expression and oncogenic functions of p21(CIP1/WAF1) in the liver. Oncogene 35, 4200–4211 (2016).

11. Niwa Y, et al. Methylation silencing of SOCS-3 promotes cell growth and migration by enhancing JAK/STAT and FAK signalings in human hepatocellular carcinoma. Oncogene 24, 6406–6417 (2005).

12. Riehle KJ, et al. Regulation of liver regeneration and hepatocarcinogenesis by suppressor of cytokine signaling 3. J Exp Med 205, 91–103 (2008).

13. Khan MGM, et al. Hepatocyte growth control by SOCS1 and SOCS3. Cytokine 121, 154733 (2019).

14. Gui Y, et al. SOCS1 controls liver regeneration by regulating HGF signaling in hepatocytes. J Hepatol 55, 1300–1308 (2011).

15. Gui Y, et al. Regulation of MET receptor tyrosine kinase signaling by suppressor of cytokine signaling 1 in hepatocellular carcinoma. Oncogene 34, 5718–5728 (2015).

16. Thorgeirsson SS, Grisham JW. Molecular pathogenesis of human hepatocellular carcinoma. Nat Genet 31, 339–346 (2002).

17. Abbas T, Dutta A. p21 in cancer: intricate networks and multiple activities. Nat Rev Cancer 9, 400–414 (2009).

18. Koster R, et al. Cytoplasmic p21 expression levels determine cisplatin resistance in human testicular cancer. J Clin Invest 120, 3594–3605 (2010).

19. Chen W, et al. Direct interaction between Nrf2 and p21(Cip1/WAF1) upregulates the Nrf2- mediated antioxidant response. Mol Cell 34, 663–673 (2009).

20. DeNicola GM, et al. Oncogene-induced Nrf2 transcription promotes ROS detoxification and tumorigenesis. Nature 475, 106–109 (2011).

21. Menegon S, Columbano A, Giordano S. The Dual Roles of NRF2 in Cancer. Trends Mol Med 22, 578–593 (2016).

22. Sitko JC, et al. SOCS3 regulates p21 expression and cell cycle arrest in response to DNA damage. Cellular signalling 20, 2221–2230 (2008).

23. Llovet JM, Montal R, Sia D, Finn RS. Molecular therapies and precision medicine for hepatocellular carcinoma. Nat Rev Clin Oncol 15, 599–616 (2018).

24. Cancer Genome Atlas Research Network. Electronic address wbe, Cancer Genome Atlas Research N. Comprehensive and Integrative Genomic Characterization of Hepatocellular Carcinoma. Cell 169, 1327–1341 e1323 (2017).

25. Liu J, et al. Integrative metabolomic characterisation identifies altered portal vein serum metabolome contributing to human hepatocellular carcinoma. Gut, (2021).

26. Fausto N, Campbell JS. Mouse models of hepatocellular carcinoma. Semin Liver Dis 30, 87–98 (2010).

27. Yan HX, et al. DNA damage-induced sustained p53 activation contributes to inflammation-associated hepatocarcinogenesis in rats. Oncogene 32, 4565–4571 (2013).

28. Hanahan D, Weinberg RA. Hallmarks of cancer: the next generation. Cell 144, 646–674 (2011).

29. Komatsu M, et al. The selective autophagy substrate p62 activates the stress responsive transcription factor Nrf2 through inactivation of Keap1. Nat Cell Biol 12, 213–223 (2010).

30. Sakurai T, et al. Hepatocyte necrosis induced by oxidative stress and IL-1 alpha release mediate carcinogen-induced compensatory proliferation and liver tumorigenesis. Cancer Cell 14, 156–165 (2008).

31. Uchida K. 4-Hydroxy-2-nonenal: a product and mediator of oxidative stress. Prog Lipid Res 42, 318–343 (2003).

32. Schaur RJ, Siems W, Bresgen N, Eckl PM. 4-Hydroxy-nonenal-A Bioactive Lipid Peroxidation Product. Biomolecules 5, 2247–2337 (2015).

33. Polonen P, et al. Nrf2 and SQSTM1/p62 jointly contribute to mesenchymal transition and invasion in glioblastoma. Oncogene 38, 7473–7490 (2019).

34. Alia M, Ramos S, Mateos R, Bravo L, Goya L. Response of the antioxidant defense system to tert-butyl hydroperoxide and hydrogen peroxide in a human hepatoma cell line (HepG2). J Biochem Mol Toxicol 19, 119–128 (2005).

35. Lu Y, Cederbaum AI. Cisplatin-induced hepatotoxicity is enhanced by elevated expression of cytochrome P450 2E1. Toxicol Sci 89, 515–523 (2006).

36. Ehedego H, et al. p21 ablation in liver enhances DNA damage, cholestasis, and carcinogenesis. Cancer Res 75, 1144–1155 (2015).

37. Marhenke S, et al. p21 promotes sustained liver regeneration and hepatocarcinogenesis in chronic cholestatic liver injury. Gut 63, 1501–1512 (2014).

38. Buitrago-Molina LE, et al. The degree of liver injury determines the role of p21 in liver regeneration and hepatocarcinogenesis in mice. Hepatology 58, 1143–1152 (2013).

39. LaBaer J, et al. New functional activities for the p21 family of CDK inhibitors. Genes & development 11, 847–862 (1997).

40. Shiraha H, Yamamoto K, Namba M. Human hepatocyte carcinogenesis (review). Int J Oncol 42, 1133–1138 (2013).

41. Milkovic L, Zarkovic N, Saso L. Controversy about pharmacological modulation of Nrf2 for cancer therapy. Redox Biol 12, 727–732 (2017).

42. Ngo HKC, Kim DH, Cha YN, Na HK, Surh YJ. Nrf2 Mutagenic Activation Drives Hepatocarcinogenesis. Cancer Res 77, 4797–4808 (2017).

43. Zhang M, et al. Nrf2 is a potential prognostic marker and promotes proliferation and invasion in human hepatocellular carcinoma. BMC cancer 15, 531 (2015).

44. Lee K, et al. The Clinicopathological and Prognostic Significance of Nrf2 and Keap1 Expression in Hepatocellular Carcinoma. Cancers (Basel) 12, (2020).

45. Guichard C, et al. Integrated analysis of somatic mutations and focal copy-number changes identifies key genes and pathways in hepatocellular carcinoma. Nat Genet 44, 694–698 (2012).

46. Zavattari P, et al. Nrf2, but not beta-catenin, mutation represents an early event in rat hepatocarcinogenesis. Hepatology 62, 851–862 (2015).

47. Umemura A, et al. p62, Upregulated during Preneoplasia, Induces Hepatocellular Carcinogenesis by Maintaining Survival of Stressed HCC-Initiating Cells. Cancer Cell 29, 935–948 (2016).

48. He F, et al. NRF2 activates growth factor genes and downstream AKT signaling to induce mouse and human hepatomegaly. J Hepatol 72, 1182–1195 (2020).

49. Parkinson EI, Hergenrother PJ. Deoxynyboquinones as NQO1-Activated Cancer Therapeutics. Acc Chem Res 48, 2715–2723 (2015).

50. Colaprico A, et al. TCGAbiolinks: an R/Bioconductor package for integrative analysis of TCGA data. Nucleic Acids Res 44, e71 (2016).

51. Love MI, Huber W, Anders S. Moderated estimation of fold change and dispersion for RNA-seq data with DESeq2. Genome Biol 15, 550 (2014).

52. Therneau TM. A Package for Survival Analysis in R. https://CRAN.R-project.org/package=survival (2020).

53. Kassambara A. Drawing Survival Curves using ‘ggplot2’. R package version 0.4.6. . http://CRAN.r-project.org/web/packages/survminer (2016).

54. Liberzon A, Subramanian A, Pinchback R, Thorvaldsdottir H, Tamayo P, Mesirov JP. Molecular signatures database (MSigDB) 3.0. Bioinformatics 27, 1739–1740 (2011).

55. Dolgalev I. msigdbr: MSigDB Gene Sets for Multiple Organisms in a Tidy Data Format., https://CRAN.R-project.org/package=msigdbr (2020).

56. Kohl M. MKmisc: Miscellaneous functions from M. Kohl_. http://www.stamats.de (2019).

57. Yu G, Wang LG, Han Y, He QY. clusterProfiler: an R package for comparing biological themes among gene clusters. OMICS 16, 284–287 (2012).

58. Foroutan M, Bhuva DD, Lyu R, Horan K, Cursons J, Davis MJ. Single sample scoring of molecular phenotypes. BMC Bioinformatics 19, 404 (2018).

59. Naugler WE, et al. Gender disparity in liver cancer due to sex differences in MyD88- dependent IL-6 production. Science 317, 121–124 (2007).

