## Supplementary figures for "SOCS3-mediated activation of p53-p21-NRF2 axis and cellular adaptation to oxidative stress in SOCS1-deficient hepatocellular carcinoma"

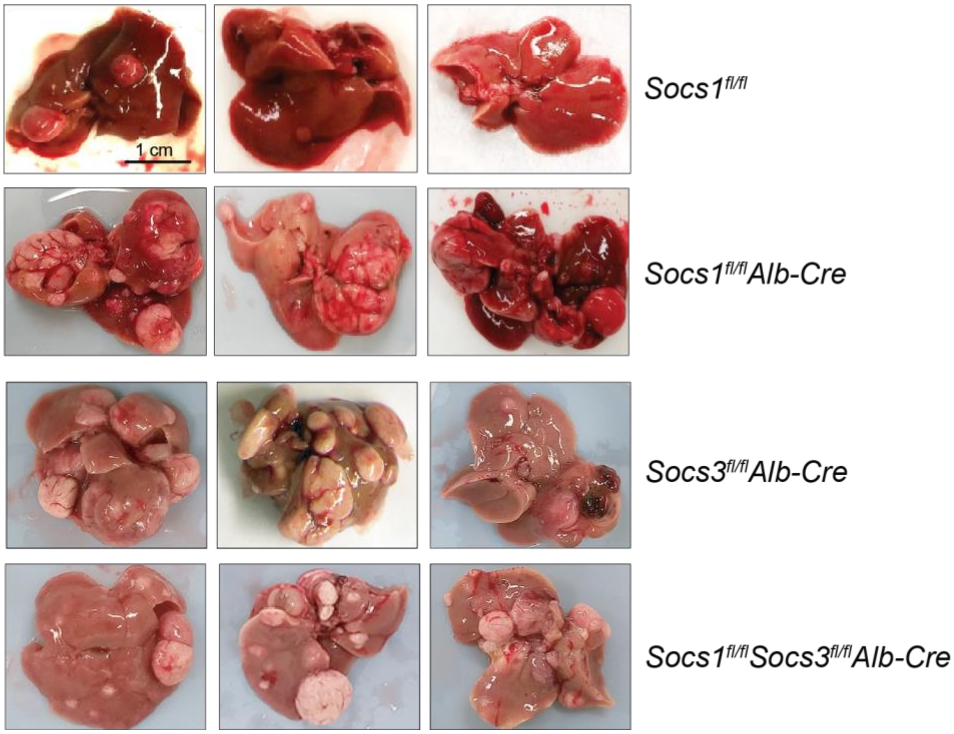


**Supplementary Figure S1.**
Increased susceptibility of SOCS1-deficient mice to HCC is mediated by SOCS3. Two weeks-old mice of the indicated genotype were treated with DEN (25 mg/Kg bodyweight, i.p.). HCC development was evaluated ten months later. Livers from three representative mice for each genotype are shown.

**Supplementary Figures**

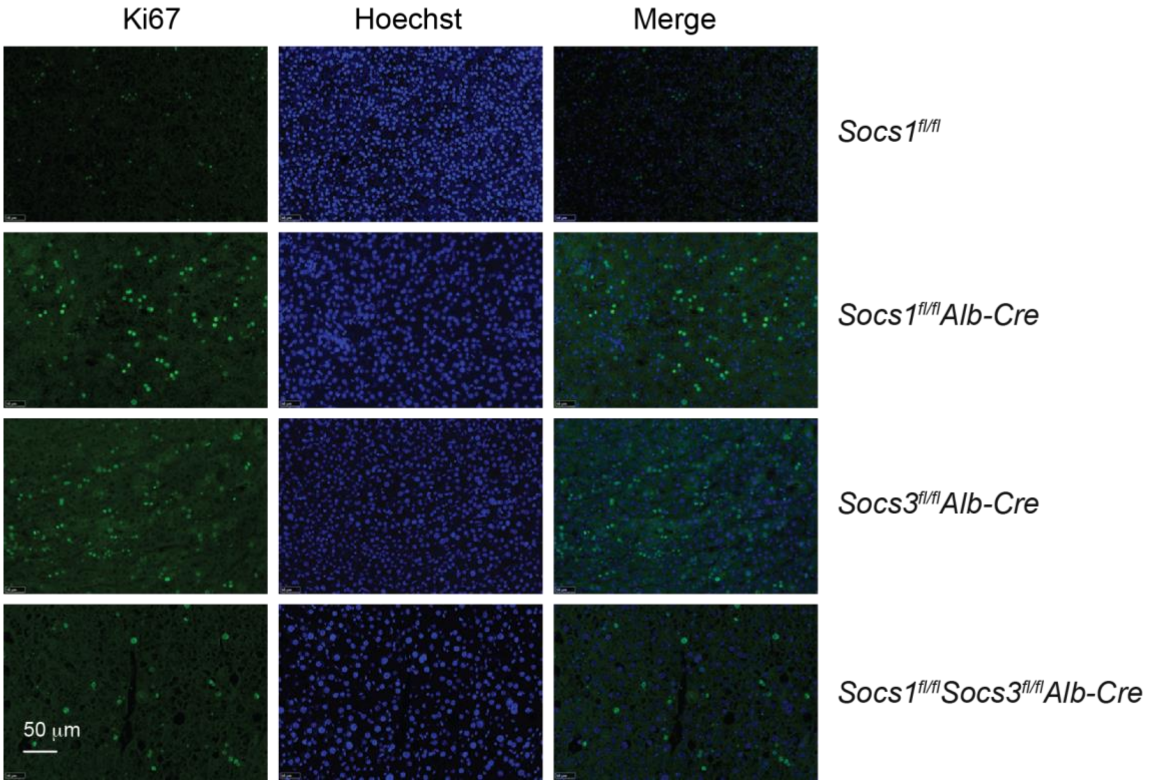


**Supplementary Figure S2.**

Increased Ki67 immunostaining in the HCC nodules of SOCS1-deficient livers is dependent on SOCS3. Liver sections from the indicated genotypes of mice bearing DEN-induced HCC were stained for Ki67. Digitally enlarged images of representative tumor nodules shown in Fig. 2D are shown.

**Supplementary Figures**

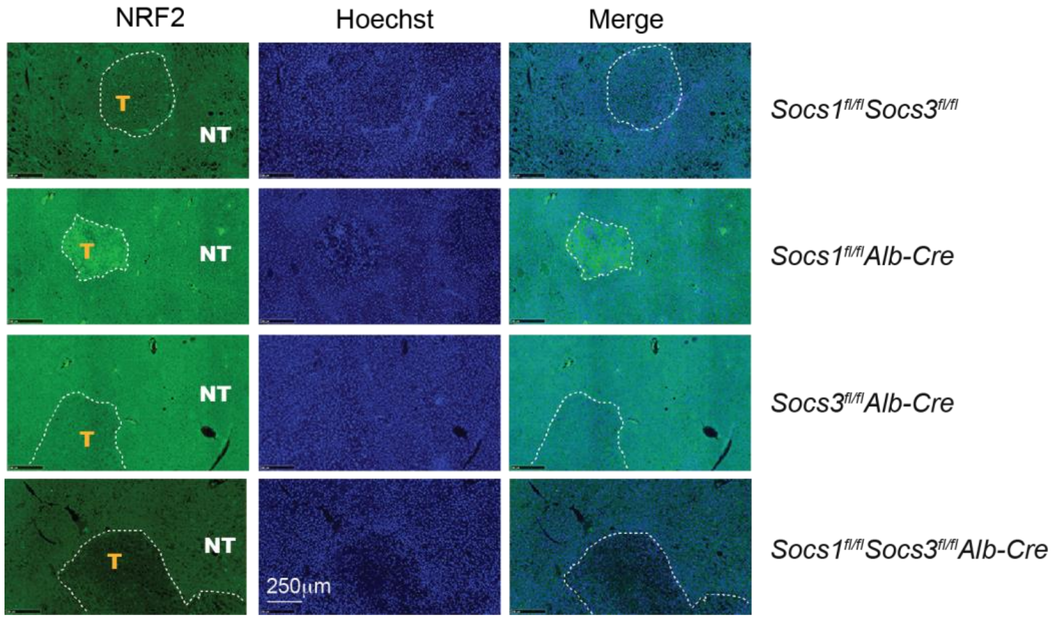


**Supplementary Figure S3.**

HCC nodules of SOCS1-deficient mice show increased NFR2 expression that is dependent on SOCS3. Liver sections from the indicated genotypes of mice bearing DEN-induced HCC were stained for NRF2. Tumor nodules (T) are demarcated from surrounding normal tissue (NT). Small tumor nodules in the knockout mice (compared to Fig. 4C) are shown for clarity.

**Supplementary Figures**

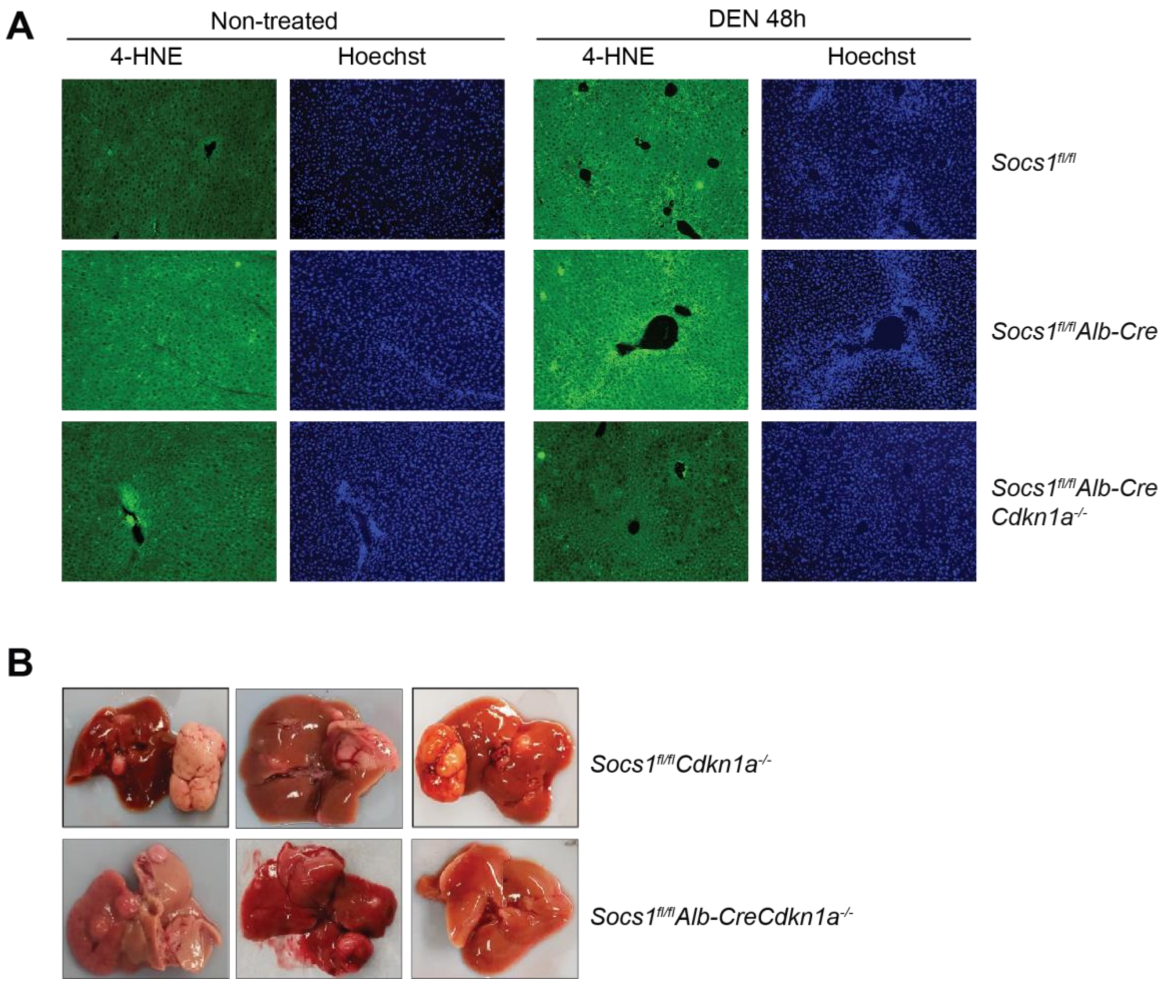


**Supplementary Figure S4.**

(A) Loss of CDKN1A reduces the ability of SOCS1-deficient hepatocytes to generate ROS needed for lipid peroxidation. Liver sections from mice of the indicated genotypes at steady-state and 48h after DEN treatment were stained for 4-HNE. Low magnification images of the sections shown in Fig. 6A are presented. Representative images at 10X magnification.

(B) CDKN1A is the crucial mediator of increased hepatocarcinogenesis in SOCS1-deficient livers. Two weeks-old mice of the indicated genotype were treated with DEN (25 mg/Kg bodyweight, i.p.). HCC development was evaluated ten months later. Livers from three representative mice for each genotype are shown.

### Supplementary Figures

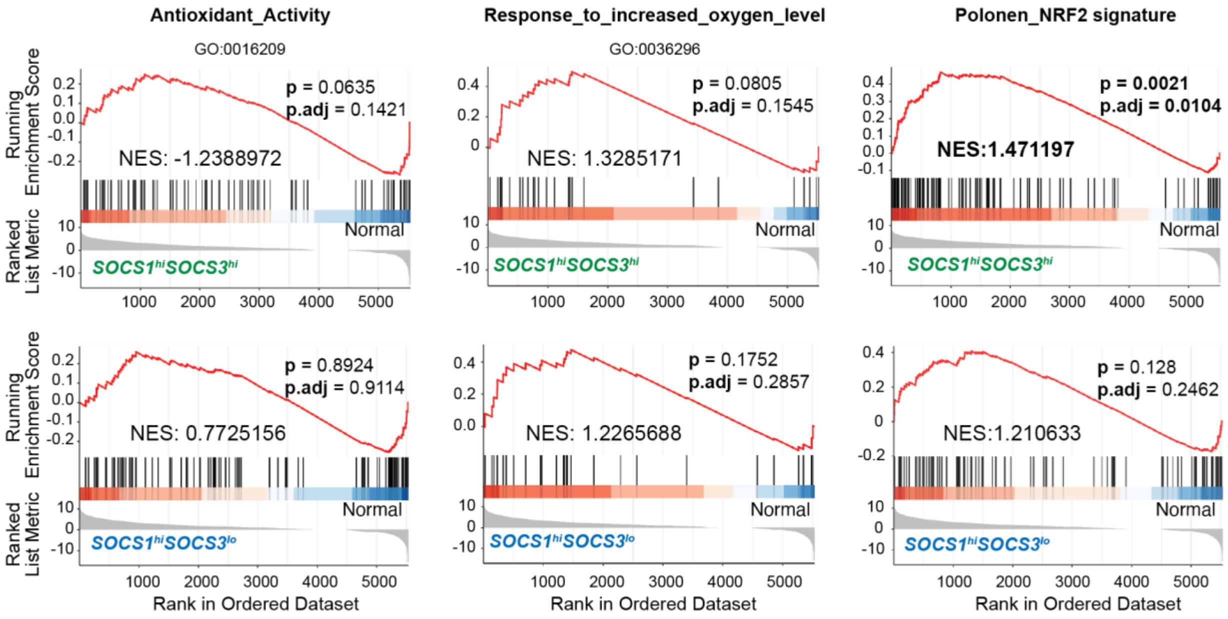


**Supplementary Figure S5.**

GSEA plots showing lack of enrichment of 'Antioxidant_Activity' and 'Response_to_increased_oxygen_level' pathway genes in *SOCS1-high/SOCS3-high* and *SOCS1-high/SOCS3-low* groups and a positive enrichment of 'NRF2 Signature' genes in *SOCS1-high/SOCS3-high* but not in *SOCS1-high/SOCS3-low* group.

Description of GO terms identified by number in Fig. 3E:

**GO:0016701**:
GO_OXIDOREDUCTASE_ACTIVITY_ACTING_ON_SINGLE_DONORS_WITH_ INCORPORATION_OF_MOLECULAR_OXYGEN.

**GO:0016634**:
GO_OXIDOREDUCTASE_ACTIVITY_ACTING_ON_THE_CH_CH_GROUP_ OF_DONORS_OXYGEN_AS_ACCEPTOR.

**GO:0016709**:
GO_OXIDOREDUCTASE_ACTIVITY_ACTING_ON_PAIRED_DONORS_WITH_ INCORPORATION_OR_REDUCTION_OF_MOLECULAR_OXYGEN_NAD_P_H_AS_ONE _ DONOR_AND_INCORPORATION_OF_ONE_ATOM_OF_OXYGEN.

**GO:00166712**:
GO_OXIDOREDUCTASE_ACTIVITY_ACTING_ON_PAIRED_DONORS_WITH_ INCORPORATION_OR_REDUCTION_OF_MOLECULAR_OXYGEN_REDUCED_FLAVIN _ OR_FLAVOPROTEIN_AS_ONE_DONOR_AND_INCORPORATION_ OF_ONE_ATOM_OF_OXYGEN.

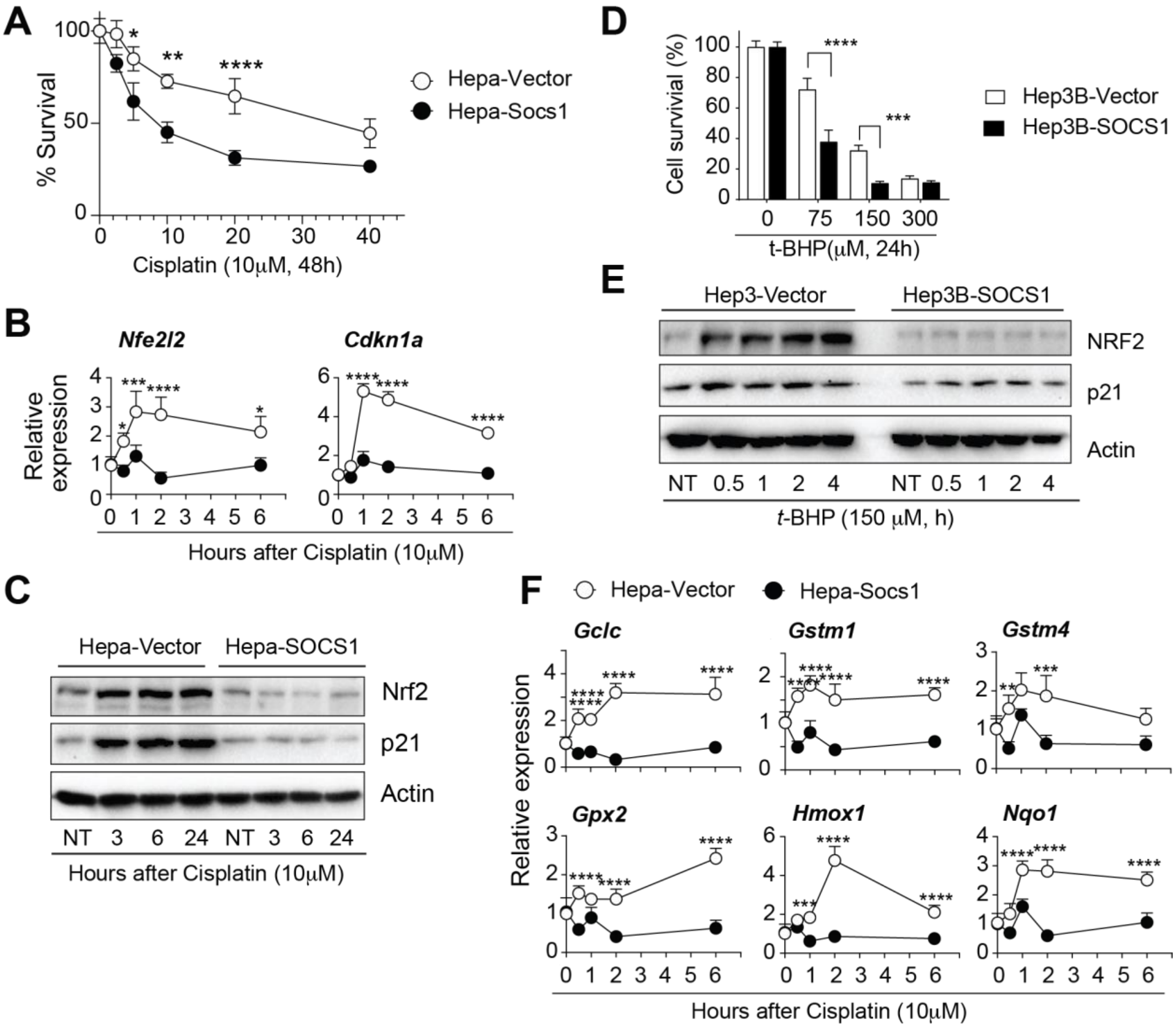


**Supplemental Figure S6.** SOCS1 sensitizes hepatocytes to oxidative stress by inhibiting NRF2 activation.
(A) Hepa-vector and Hepa-SOCS1 were treated with cisplatin for 48h and cell survival was evaluated (n=3).
(B,C) Hepa-vector and Hepa-SOCS1 cells were treated with cisplatin and the induction of *Nfe2l2* and *Cdkn1a* genes (B, n=4) and protein levels of NRF2 and p21 (C, data representative of two experiments) were evaluated at the indicated time points.
(D) Hep3B human HCC cells expressing control vector (Hep3B-vector) or SOCS1 (Hep3B-SOCS1) were treated with *t*-BHP for 24h and cell survival was evaluated (n=3).
(E) Hep3B-vector and Hep3B-SOCS1 cells treated with t-BHP and NRF2 and p21 protein levels were evaluated at the indicated time points. Representative data from two experiments.
(F) Hepa-vector and Hepa-SOCS1 cells were treated with cisplatin and the induction of NRF2-induced antioxidant genes was measured by qRT-PCR (n=4).
Statistics: Mean + SEM; Two-way ANOVA with Tukey's multiple comparison test: *$p<0.05$, **$p<0.01$, ***$p<0.001$, ****$p<0.0001$.
