## Supplementary Tables for "SOCS3-mediated activation of p53-p21-NRF2 axis and cellular adaptation to oxidative stress in SOCS1-deficient hepatocellular carcinoma"

**Supplemental Table S1:** List of Mouse strains used in this study.

| Strain | Citation | Supplier |
|---|---|---|
| *Socs1$^{fl/fl}$* | PMID: 18322180 | Dr. Akihiko Yoshimura, Tokyo |
| *Socs3$^{fl/fl}$* | PMID: 12754507 | Jax mice, Stock # 010944 |
| *Alb$^{Cre}$* | PMID: 9867845 | Jax mice, Stock # 003574 |
| *Cdkn1a$^{-/-}$* | PMID: 7566157 | Jax mice, Stock # 003263 |
| *Tp53$^{-/-}$* | PMID: 7922305 | Jax mice, Stock # 002101 |
| *Socs1$^{fl/fl}$ Alb$^{Cre}$* | PMID: 26725321 | Previously generated |
| *Socs3$^{fl/fl}$ Alb$^{Cre}$* | PMID: 32807134 | Previously generated |
| *Socs$^{fl/fl}$ Socs3$^{fl/fl}$* | PMID: 32807134 | Previously generated |
| *Socs$^{fl/fl}$ Socs3$^{fl/fl}$ Alb$^{Cre}$* | | Generated for this study |
| *Cdkn1a$^{-/-}$Socs1$^{fl/fl}$ Alb$^{Cre}$* | | Generated for this study |
| *Tp53$^{-/-}$Socs1$^{fl/fl}$ Alb$^{Cre}$* | | Generated for this study |

**Supplemental Table S2:** List of reagents used in this study.

| Name | Supplier | Cat # |
|---|---|---|
| *tert*-butyl hydroperoxide (*t*-BHP) | Sigma-Aldrich | 458139 |
| CellROX Green | Molecular Probes (Thermo-Fisher) | C10444 |
| CellROX Deep Red | Molecular Probes (Thermo-Fisher) | C10422 |
| Cisplatin (*cis*-diamino-dichloroplatinum; *cis*-Pt) | Sigma-Aldrich | C2210000 |
| Diethylnitrosamine (DEN) | Sigma-Aldrich | N0258 |
| Glutathione Colorimetric Detection Kit | Invitrogen | EIAGSHC |
| Lipofectamine RNAiMAX | Invitrogen (Thermo-Fisher) | Cat# 13778-075 |
| RNeasy Mini Kit | Qiagen | 74104 |
| RNAlater® | Qiagen | 76106 |
| WST-8 | Dojindo Molecular Technologies | CK04 |
| QIAzol® Lysis Reagent | Qiagen | 79306 |

**Supplementary Tables**

**Supplemental Table S3:** List of software used.

| Software name | Manufacturer | Version |
|---|---|---|
| Adobe Photoshop/Illustrator | Adobe Inc. | CC |
| CFX manager | Bio-Rad Laboratories, Inc | 3.1 |
| Image Lab™ | Bio-Rad Laboratories, Inc | 6 |
| NDP.view2 | Hamamatsu Photonics K.K | |
| Prism | GraphPad | 9 |

**Supplemental Table S4:** List of antibodies used in this study.

| Name | Citation | Supplier | Cat no. | Clone no. |
|---|---|---|---|---|
| **Immunofluorescence** | | | | |
| Rabbit anti- Ki67 | PMID: 20152769 | Cell Signaling Technology | #9129 | D3B5 |
| Rabbit anti-NRF2 (mAb) | PMID: 32057361 | Cell Signaling Technology | #12721 | D1Z9C |
| Mouse anti-4-hydroxy-nonenol (4-HNE) | PMID: 30878614 | abcam | Ab48506 | HNEJ-2 |
| **Western blot** | | | | |
| Rabbit-anti p21 | PMID: 20959475 | Santa Cruz Biotechnology | SC-471 | Polyclonal |
| Mouse-anti p21 | PMID: 32499530 | Santa Cruz Biotechnology | SC-817 | Not given |
| Rabbit anti-NRF2 (mAb) | PMID: 19417020 | Cell Signaling Technology | #12721 | D1Z9C |
| Rabbit anti-Nrf2 | PMID: 30896860 | abcam | ab156883 | Polyclonal |
| Rabbit anti β-Actin | PMID: 19933848 | Cell Signaling Technology | #4970 | 13E5 |
| Mouse anti-p53 | PMID: 26725321 | Cell Signaling Technology | #2524 | 1C12 |
| Rabbit anti-Phospho p53 (Ser15) | PMID: 26725321 | Cell Signaling Technology | #9284 | Polyclonal |
| Rabbit anti-Stat1 (p84/p91) | PMID: 26725321 | Santa Cruz Biotechnology | SC-592 | M-22 |
| Rabbit anti-Phospho Stat1 (Y701) | PMID: 26725321 | Cell Signaling Technology | #9171 | Polyclonal |
| Rabbit anti-Stat3 | PMID: 26725321 | Santa Cruz Biotechnology | SC-483 | Polyclonal |
| Rabbit anti-Phospho Stat3 (Y705) | PMID: 26725321 | Cell Signaling Technology | #9131 | Polyclonal |

### Supplementary Tables

**Supplemental Table S5:** List of cell lines used in this study.

| Name | Citation | Supplier | Cat no. | Authentication test method |
|---|---|---|---|---|
| Hepa1-6 (Hepa) | PMID: 3431441 | ATCC | CRL-1830 | STR profiling (according to the product sheet) |
| Hepa-SOCS1 | PMID: 21703184 | Generated in our previous study | | Derived from Hepa 1-6 |
| Hep3B | PMID: 6248960 | ATCC | HB-8064 | STR (according to the product sheet) |
| Hep3B-SOCS1 | PMID: 25728680 | Generated in our previous study | | Derived from Hep3B |

***Note:*** The two lines Hepa1-6 and Hep3B do not appear in the misidentified cell line list maintained by the International Cell Line Authentication Committee (ICLAC). STR, short-tandem repeat profiling.

**Supplemental Table S6:** List of siRNA reagents used.

| siRNA | Source | Catalogue # |
|---|---|---|
| si-Scramble | Dharmacon | Cat# D-001210-01-05 |
| si-p21-V1 | SigmaAldrich | SASI Mm01 00086762 (sequence start at 511) |
| si-p21-V2 | SigmaAldrich | SASI Mm01 00086763 (sequence start at 516) |
| si-p21-V3 | SigmaAldrich | SASI Mm02 00306392 (sequence start at 536) |

### Supplementary Tables

**Supplemental Table S7:** List of RT-qPCR primers used in this study.

| Gene Name | Gene ID | Forward Sequence (5'-3') | Reverse Sequence (5'-3') | Amplicon size (bp) |
|---|---|---|---|---|
| *36B4 (Rplp0)* | NM_007475.5 | TCTGGAGGGTGTCCGCAA | CTTGACCTTTTCAGTAAGTGG | 154 |
| *Cdkn1a* | NM_007669.5 | TTCTATCACTCCAAGCGCAG | CAGGCAGCGTATATACAGGAG | 185 |
| *Gadd45a* | NM_007836.1 | AGTCAGCGCACCATTACG | TGAGGGTGAAATGGATCTGC | 139 |
| *Gclc* | NM_010295.2 | ACCATCACTTCATTCCCCAG | TTCTTGTTAGAGTACCGAAGCG | 148 |
| *Gpx1* | NM_008160.6 | GACTACACCGAGATGAACGATC | TCTCACCATTCACTTCGCAC | 199 |
| *Gpx2* | NM_030677.2 | GTAGTTCTCGGCTTCCCTTG | GGTAAGACTAAAGGTGGGCTG | 123 |
| *Gstm1* | NM_001374678.1 | CTATGATACTGGGATACTGGAACG | ACTTCTCATTCAGCCACTGG | 147 |
| *Gstm4* | NM_026764.3 | TGAAGGTGGAATACTTGGAGC | GGGTTCGAATATAAGGTGCAGG | 149 |
| *Hmox1* | NM_010442.2 | ACAGAGGAACACAAAGACCAG | GTGTCTGGGATGAGCTAGTG | 136 |
| *Keap1* | NM_001110306.1 | CTCCGCAGAATGTTACTATCCAG | ACACTGTTCAACTGGTCCTG | 147 |
| *Mdm2* | NM_010786.4 | TGGCGTAAGTGAGCATTCTG | CTGTATCGCTTTCTCCTGTCTG | 188 |
| *Nfe2l2* | NM_010902.4 | TCCCATTTGTAGATGACCATGAG | CCATGTCCTGCTCTATGCTG | 150 |
| *Nqo1* | NM_008706.5 | TGAAGAAGAGAGGATGGGAGG | GATGACTCGGAAGGATACTGAAAG | 130 |
| *Sesn1* | NM_001162908.1 | GGACGAGGAACTTGGAATCAG | CAAAGGAGTCTGCAAATAACGC | 131 |
| *Sesn2* | NM_144907.1 | CAGCCTCACCTATAACACCATC | CTCGCCGTAATCATAGTCATCG | 124 |
| *Socs1* | NM_001271603.1 | TGGTTGTAGCAGCTTGTGTCTGG | CCTGGTTTGTGCAAAGATACTGGG | 92 |
| *Socs3* | NM_007707.3 | GAGATTTCGCTTCGGGACTAG | TTTGGAGCTGAAGGTCTTGAG | 193 |
| *Tp53* | NM_011640.3 | AGTTCATTGGGACCATCCTG | GCTGATATCCGACTGTGACTC | 149 |
